# High multiplex, digital spatial profiling of proteins and RNA in fixed tissue using genomic detection methods

**DOI:** 10.1101/559021

**Authors:** Christopher R. Merritt, Giang T. Ong, Sarah Church, Kristi Barker, Gary Geiss, Margaret Hoang, Jaemyeong Jung, Yan Liang, Jill McKay-Fleisch, Karen Nguyen, Kristina Sorg, Isaac Sprague, Charles Warren, Sarah Warren, Zoey Zhou, Daniel R. Zollinger, Dwayne L. Dunaway, Gordon B. Mills, Joseph M. Beechem

## Abstract

We have developed Digital Spatial Profiling (DSP), a non-destructive method for high-plex spatial profiling of proteins and RNA, using oligonucleotide detection technologies with unlimited multiplexing capability. The key breakthroughs underlying DSP are threefold: (1) multiplexed readout of proteins/RNA using oligo-tags; (2) oligo-tags attached to affinity reagents (antibodies/RNA probes) through a photocleavable (PC) linker; (3) photocleaving light projected onto the tissue sample to release PC-oligos in any spatial pattern. Here we show precise analyte reproducibility, validation, and cellular resolution using DSP. We also demonstrate biological proof-of-concept using lymphoid, colorectal tumor, and autoimmune tissue as models to profile immune cell populations, stroma, and cancer cells to identify factors specific for the diseased microenvironment. DSP utilizes the unlimited multiplexing capability of modern genomic approaches, while simultaneously providing spatial context of protein and RNA to examine biological questions based on analyte location and distribution.

## INTRODUCTION

Understanding the spatial distribution of protein and RNA has led to important discoveries for defining tissue pathology and discovering biomarkers that predict patient response to therapy ^1-6^. However, there remains much to learn about better targets for drugs, predictors of response to therapy, and biological mechanisms of action based on protein and RNA localization within tissues. Immunohistochemistry (IHC) and RNA *in situ* hybridization (ISH) have traditionally been used to assess the spatial distribution of proteins and RNAs in fixed tissue samples. However, these techniques are limited in their multiplexing and quantitation of markers by the optical nature of chromogenic and fluorescent detection chemistries. The importance of investigating the expression of high numbers of markers in a spatially resolved manner has motivated the development of several new methodologies to increase multiplexing capabilities. These higher-plex methodologies tend to fall into two broad classes: 1) *sequential spatial analyses*, which examine tissues over several reagent processing cycles across a relatively small number fluorescent channels, and 2) *simultaneous spatial analyses*, which utilize readouts that do not involve light detection, and are thus not constrained by the number of fluorescent probes that can be used within the light spectrum. The most widely used commercial methods for sequential spatial analysis are Perkin Elmer’s multispectral imaging system^7^ and RNAscope (ACD)^8^ technologies for antibody-based protein detection and oligonucleotide-based RNA detection, respectively. Both technologies utilize tyramide signal amplification (TSA) to amplify signal, and therefore sensitivity. Since these approaches are limited by the ability to resolve independent fluorophores, the maximum plexing capabilities of these technologies is typically around 4-8 markers per section, depending on the detection microscopes capability. Codetection by Indexing/CODEX (Akoya)^9^ and InsituPlex (Ultivue)^10^ are sequential analysis methodologies developed for protein analysis that increase target plex through labeling antibodies with unique oligonucleotides. These oligonucleotide tags can be read using fluorescent detection during amplification of the probe sequence or through the cyclical hybridization of fluorescently-labeled reverse complement sequences. Multiplexed error-robust fluorescence *in situ* hybridization/MERFISH^11^ and fluorescent *in situ* RNA sequencing/FISSEQ (ReadCoor)^12^ are two technologies under development that rely on fluorescent detection sequencing approaches for determining RNA probe expression. A general limitation for technologies that use sequential spatial analysis is the impact of multiple hybridizations on a single section, which typically decreases sample throughput, and increases the time to results.

Recent developments in simultaneous spatial analysis have been driven by the utilization of mass spectrometry (MS) without affinity reagents or further enhanced approaches using isotope metal labeled antibodies that can be visualized with MS. Techniques that do not rely on affinity reagents, such as Matrix-assisted laser desorption/ionization (MALDI) imaging mass spectrometry (IMS)^13^ have the ability to profile very high-plex numbers of peptides and proteins in a spatial manner. MALDI-IMS uses aromatic molecules in a matrix that forms crystals that can be absorbed and ionized by UV laser and subsequently measured by mass spectrometry to give protein expression for hundreds of proteins.^14^ The two leading technologies using affinity reagents coupled with mass-spectrometry (MS) are Imaging Mass Cytometry/IMC (Fluidigm)^15^ and Multiplexed Ion Beam Imaging/MIBI (IonPath).^16^ MS readout of metal isotope labeled antibodies has the ability to extended to ~100-plex targets, but is limited primarily by the number of metal isotopes that can be conjugated to the affinity reagents and the relatively high cost of MS instrumentation. Another technology using simultaneous spatial analysis, Spatial Transcriptomics^17^ performs simultaneous spatial analysis of RNA on non-fixed tissue through sequencing readout of RNA transcripts captured on a sample-independent predefined grid. In general, most of these approaches are limited by the maximum target number that can be detected, and the ability to analyze only one analyte type (protein or RNA). The benefit of nearly all of these approaches (except spatial transcriptomics), is that high resolution images are output, which are very similar to those produced with traditional approaches, and enable sophisticated analyses with digital imaging process software.^18, 19^

The GeoMx™ Digital Spatial Profiling (DSP) system (NanoString Technologies, Inc.) uses a novel tissue-sampling, region of interest (ROI) approach for simultaneous spatial analysis of protein and RNA. With this approach, small DNA oligonucleotide tags are attached to the affinity reagents (antibodies or RNA hybridization probes) through a photocleavable linker allowing precise spatially resolved release in ROIs (photocleaved analyte detection oligo, or PC-oligo). The spatially resolved PC-oligo is released into an aqueous buffer layer over the tissue slice and aspirated into an on-instrument microtiter plate using a microcapillary that is centered above the photocleaving illumination area. The photocleaving light is projected onto the tissue sample using automated, programmable, digital-micromirror device (DMD), allowing complete flexibility in the spatial-domain of the high-plex profiling. ROIs can either be spatially restricted or composed of multiple areas across a slide to encompass specific cell types. The PC-oligo approach is compatible with standard immunofluorescence approaches allowing guided selection of regions or cells of interest. These PC-oligos are finally digitally quantified through the nCounter System (NanoString Technologies, Inc) or a Next Generation Sequencing (NGS) platform (*e.g.*, Illumina MiSeq or NextSeq).

The GeoMX DSP system has been developed with hardware, software, and chemistry to allow for up to 800-plex profiling of mRNA or protein using an optical-barcode readout^20, 21^, and potential for unlimited multiplexing using the NGS readout. Full results from 10 to 20 slides (depending on number of ROIs) can be accomplished within 1.5 to 2.5 days. The system is highly sensitive, simple to run, and produces direct, digital data for quantitative downstream analysis. Importantly, the technology is non-destructive, allowing multiple cycles of high-plex profiling on the same tissue section or subsequent DNA sequencing on the same section. With a prototype DSP instrument, we demonstrate up to 44-plex protein characterization of a number of unique types of ROIs including geometric, gridded, segmented, rare cell, and contour population profiling. Spatially-resolved RNA expression, demonstrated with 96 target genes with using nCounter System detection and at >900-plex using NGS detection, allows study of transcriptional and post-transcriptional regulation. Applying different ROI selection methods in DSP we examined human formalin-fixed paraffin-embedded (FFPE) tonsil, colorectal tumors and inflammatory bowel disease (IBD) tissue to show the ability to decipher distinct heterogeneity of immune biology in different compartments of the tissue, tumor microenvironment (TME), as well as autoimmune disease microenvironment. We also demonstrate how DSP can be used to discover immune markers spatially located within tissue as prognostic and predictive potential for clinical decisions.

## RESULTS

### Digital Spatial Profiling (DSP) platform overview

The DSP platform quantifies the abundance of protein or RNA by counting unique indexing oligos assigned to each target of interest (**Fig. 1a**). Indexing oligos are covalently attached to primary antibodies or to mRNA hybridization probes with a UV-photocleavable linker.

**Fig. 1:**
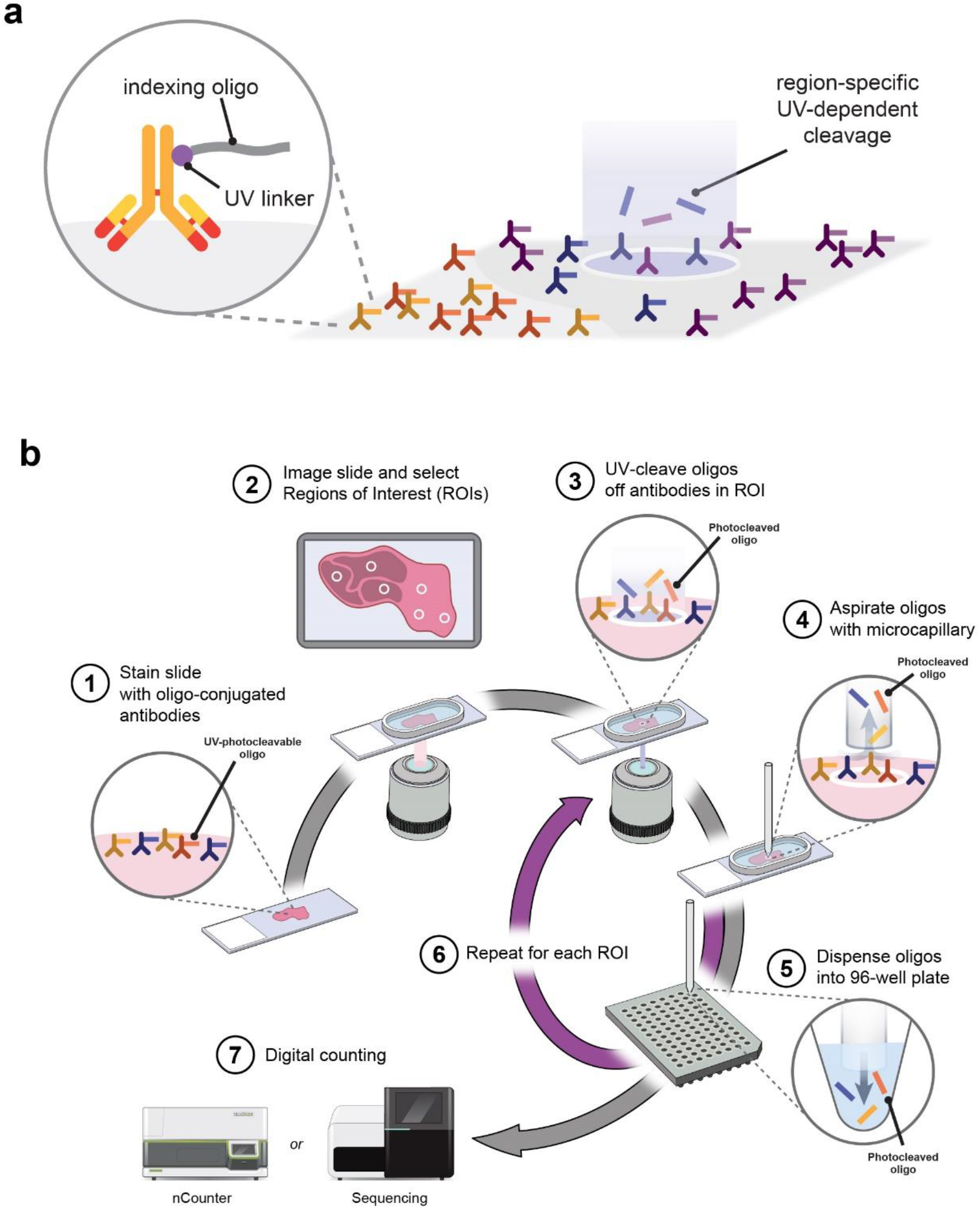
Overview of DSP workflow. **a**, DSP concept. Affinity reagents (e.g. antibodies or RNA probes) are covalently linked to a DNA indexing oligo with a UV-cleavable linker (PC-oligo). These affinity reagents are used to stain tissue sections and focused UV light can precisely liberate indexing oligos from any region of interest. These oligos can then be collected and digitally quantified. **b**, Overview of localized protein profiling workflow on the DSP instrument. (1) Process tissue section: FFPE slide-mounted tissue is subjected to a standard antigen retrieval protocol, followed by incubation with fluorescent imaging reagents (specific antibodies or biological stains and dyes) and a cocktail of primary antibodies conjugated to PC-oligos. (2) View and select ROIs: DSP scans 20x image of up to 4-channel fluorescence. Fluorescent imaging reagents establish overall architecture of tissue (e.g., PanCK tumor cells, SYTO83 nuclei and CD45 immune cells). ROIs are drawn or automatically places based on fluorescence for multiplex profiling. (3) Localized PC-oligo release: Oligos from the selected ROI are released upon localized exposure to UV light. (4-5) Oligo collection: Photocleaved oligos that have been released into aqueous liquid above the slide are collected via a microcapillary tube and deposited in a 96-well plate for subsequent quantitation. (6) Repeat: Localized UV exposure and PC-oligo collection (Steps 3-5) are repeated for each ROI, with extensive washing between each cycle. (7) Digital or NGS count: Spatially resolved pools of photocleaved oligos are either: hybridized to NanoString’s fluorescent barcodes, enabling the digital counting of up to ~1 million binding events per ROI using the standard NanoString nCounter Analysis system or by NGS quantification: the entire plate is pooled into a single tube, purified, QC’d and sequenced. NGS reads are processed into digital counts and mapped back to each ROI, generating a map of protein and transcript activity within the tissue architecture.

The profiling process begins with a slide-mounted FFPE tissue section that undergoes antigen retrieval procedures or mild Proteinase K digestion prior to incubation with a cocktail of photocleavable-oligo-labeled primary antibodies or mRNA-binding probes, respectively. The same section, or a serial section, is also stained with up to four visible imaging reagents to identify tissue features and biomarkers of interest. Simultaneously to the imaging reagents, oligo-conjugated affinity reagents are applied to the sample, resulting in a single-step for reagent application (**Fig. 1b**). Once the incubation is complete, slides are loaded onto the DSP instrument and each sample is scanned to produce a digital image of the tissue morphology based on the fluorescent markers. This whole slide scanned image will be the “guide” to select ROIs for profiling the high-plex PC-oligo conjugated analytes. ROIs of any shape, including non-contiguous areas, can be selected for automated molecular profiling. A programmable digital micromirror device (DMD) or dual-DMD (DDMD) directs UV light to precisely illuminate the ROI and cleave PC-oligos in a region-specific manner. DMDs are small semiconductor chips (Texas Instruments) containing millions of programmable mirrors. Any image file can be directly downloaded to a DMD and the mirrors automatically adjust in order to take a non-spatially resolved light source and, via reflection from the semiconductor mirrored surface, generate a reflected black-white (*i.e.*, on-off) replica of the image onto the tissue surface. The DMD will autoconfigure to match the exact spatial pattern from binarized visible-wavelength images or externally generated image-files of each tissue section (*e.g.,* ROIs selected on images from ImageJ or other digital image processing platform), allowing for single-step molecular profiling of the completely unique morphology and distribution of cell types in each tissue slice.

The released indexing oligos are collected via microcapillary aspiration, dispensed into a microplate, and digitally counted using the single-molecule counting nCounter System or analyzed using NGS. On the prototype instrument, the entire workflow, from tissue preparation to data output, could be completed in 1.5 to 2.5 days. (8-12 hours sample prep/antibody incubation (**Fig. 1a**: Step 1), 4-8 hours on microscope (**Fig. 1a**: Steps 3-6), 4 to 12 (same-day protocol) or 24 hours (overnight protocol) hybridization/quantification time (**Fig. 1a**: Step 7). Most steps can be combined to process multiple samples in parallel, allowing for the collection of ~384 ROIs from one or multiple specimens within one week.

### Antibody specificity following oligo conjugation

FFPE-qualified antibodies were conjugated with a PC-oligo compatible with nCounter System gene expression or NGS platforms (**Fig. 2a**). Oligos with unique, detectable sequences were covalently attached to antibodies through amine or thiol conjugation. Antibodies were tested to validate their staining specificity before and after conjugation by IHC (**Fig. 2b, Supplementary Table 1**). Standard IHC was performed on selected FFPE tissue samples or cell pellets (screened to be positive for the antibody target) on serial sections with conjugated and unconjugated antibodies. Oligo conjugation typically maintains specificity but leads to lower IHC staining intensity, part of which is likely due to blockage of some secondary antibody binding sites used during amplification in the IHC approach. This should not alter the affinity with DSP as no secondary antibodies are used. Furthermore, this modest loss of staining intensity is not problematic for the DSP readout due to the intrinsic sensitivity of the nCounter System and NGS analysis. To obtain similar staining intensity for comparison, conjugated antibodies were stained with higher concentrations of antibody relative to their unconjugated counterparts. After conjugation, in the human tonsil CD3, a marker for T cells, expression retained the same membranous pattern with strongest staining outside of the germinal centers in T cell-enriched areas (**Fig. 2b**). CD68, a marker for macrophages, expression patterns remained the same pre-and post-conjugation with strong staining both inside and outside the germinal centers localized to both the cellular membrane and cytoplasm (**Fig. 2b**). Similarly, B7-H3 and Pan-Cytokeratin (Pan-CK) retained similar staining patters pre-and post-conjugation (**Fig 2b**). In rare cases, we observed altered staining patterns following oligo conjugation, these antibodies were not used in DSP experiments. Alternate antibodies to the same target were then sought.

**Fig. 2:**
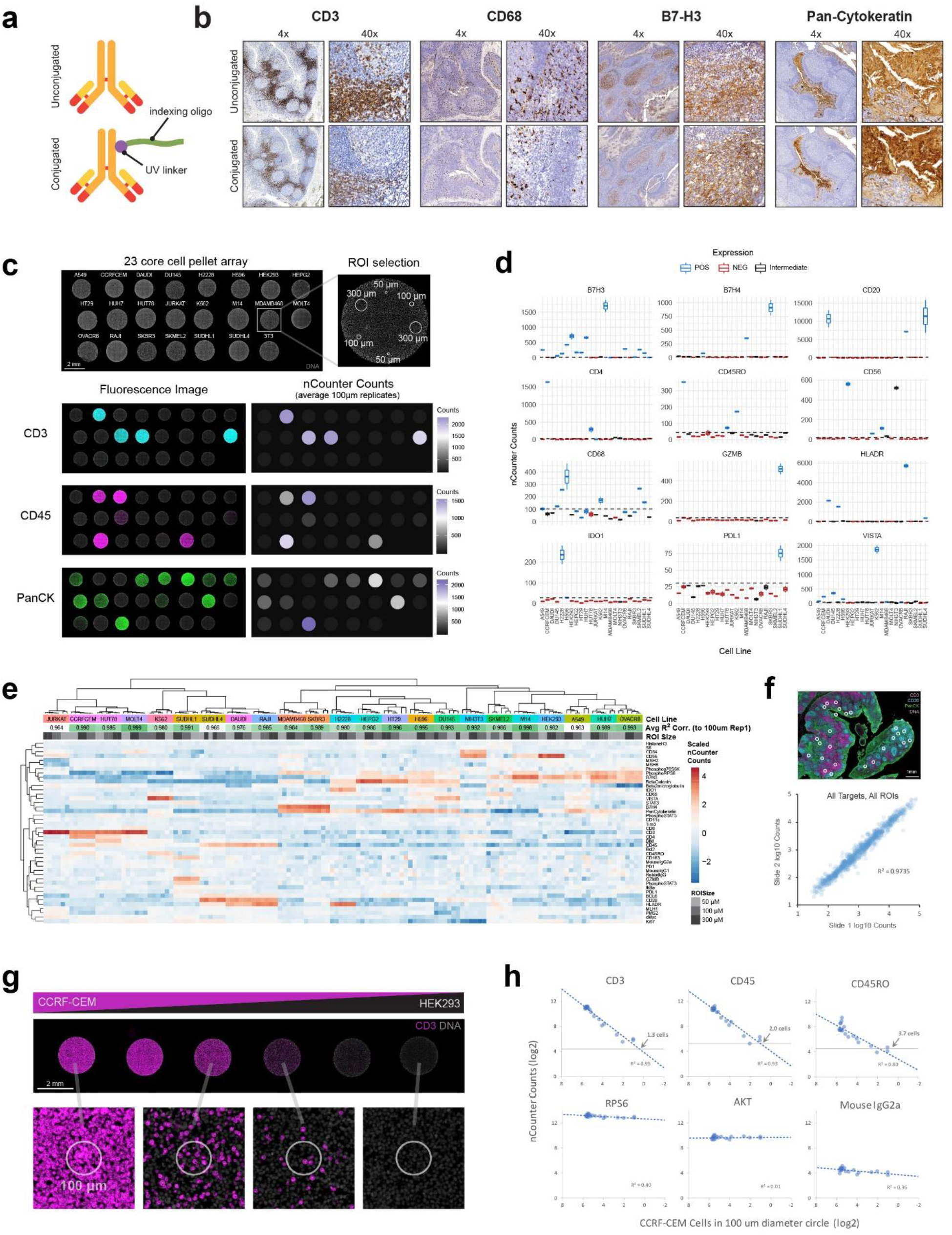
Validation of DSP with oligo-conjugated antibodies. **a**, Oligo labeled antibody schematic. Unconjugated antibodies are covalently conjugated to countable indexing oligos with a UV-cleavable moiety. **b**, Comparison of unconjugated and oligo-conjugated CD3, CD68, B7-H3, and Pan-Cytokeratin antibody staining specificity on tonsil with DAB IHC. IHC performed with unconjugated antibodies at 0.25 µg/ml, 1 µg/ml, 0.25 µg/ml, and 0.25 µg/ml, respectively. IHC performed with conjugated antibodies at 1 µg/ml, 4 µg/ml, 4 µg/ml, 1 µg/ml, respectively. **c**, 23 core cell pellet tissue microarray used for antibody validation stained with SYTO13 nuclear dye to highlight different cores. Two replicate ROIs across three profiled sizes, 50 µm, 100 µm, and 300 µm diameter circles. Fluorescently labeled antibodies for CD3, CD45, and Pan-CK were included with a mix of 44 oligo-conjugated antibodies for reference. Average nCounter counts from DSP profiling for the 100 µm ROIs displayed in a heatmaps next to the single channel fluorescent images for each target. **d**, Box plots of replicate 100 µm ROIs across all cell lines for 12 additional targets (additional targets shown in **Supplementary Fig. 1**). Positive (POS), negative (NEG) and intermediate protein expression was predicted from data on transcript abundance across cell lines, or from data previously published by others. The LOD of each target was calculated as the mean + 3x standard deviation of all NEG cell lines. **e**, Unsupervised clustering of scaled data across all ROIs. Data was scaled to characterize general expression patterns for each ROI, while considering variable ROI sizes. Average R^2^ correlations across the 6 ROIs for each cell line were determined by comparing the first replicate of the 100 µm ROI against the 5 other ROIs. These 5 independent R^2^ values were average to obtain the values shown across the heatmap annotation row [Avg R^2^ Corr. (to 100um Rep1)]. **f**, Reproducibility of counts between two tonsil serial sections. One serial section with 25 selected 200 µm circular ROIs shown. Fluorescently conjugated antibodies for CD3 (pink), CD20 (cyan), and Pan-CK (green) were used for visualization and alignment ROIs on serial sections. Correlation between normalized counts for all antibodies and all ROIs was determined. **g**, Cell pellet TMA with variable percentages of two cell lines with differential expression of CD3 (magenta). CCRF-CEM titrated into a background of HEK293 cells with approximate amounts of CCRF-CEM cells at 100%, 95%, 70%, 30%, 5%, 0%. 100 µm circles were selected across cell pellet titrations. 100%, 70%, 30%, and 0% representative ROIs shown. **h**, nCounter counts plotted against the number of CCRF-CEM cells (CD3+ cells) in each ROI for 3 targets expressed exclusively in CCRF-CEM (CD3, CD45, CD45RO), 2 targets expressed in both cell lines (RPS6 and AKT), and 1 negative control antibody (Mouse IgG2a). The LOD of each target was calculated as the mean + 3x standard deviation of all ROIs that contained no CCRF-CEM cells. Cell labels on each graph is the estimated number of cells needed for detection of the antibody.

### Validation of oligo-conjugated antibodies on the DSP system

Validation of the DSP methodology was first performed on well characterized cell lines with stable expression of most protein targets of interest. Antibodies for 44 immuno-oncology proteins (and controls) were selected and conjugated with PC-oligos for these studies (**Supplementary Table 1**). Positive and negative control cell lines were identified for targets of interest by performing multiplex gene expression analysis with 770 genes covering cancer-associated canonical pathways and different immune cell types or based on published literature. The selected cell lines were used to construct FFPE cell pellet tissue microarrays (TMAs) to quantify conjugated antibody specificity and sensitivity on the DSP system (**Fig. 2c**). One mouse cell line, NIH3T3, was included as a negative control for human antibodies, but tended to give unpredictable positive signal for many antibodies in the cocktail, and was eliminated from most analyses. This high background is likely due to high similarity of sequence between species, as we observed antigen specificity of the same antibodies in mouse tissue (*data not shown*). Whole slide imaging of the slide with nuclear staining (SYTO 13) showed the spatial distribution of the 23 TMA cores. Fluorophore conjugated antibodies for CD3, CD45, and Pan-CK were included with a mix of 44 oligo-conjugated antibodies, stained with a modified IHC protocol compatible with oligo-conjugated antibodies, and profiled on a prototype DSP instrument (see Methods for details). Six ROIs were profiled for each cell pellet: two replicate ROIs across three ROI sizes, 50 µm, 100 µm, and 300 µ m. In total, 5934 datapoints were obtained for this single DSP experiment (23 cell pellet cores × 6 ROIs × 43 oligo-conjugated antibodies).

Digital counts from the nCounter System were compared to the fluorescent images obtained on the same section for fluorescently labeled CD3, CD45, and Pan-CK antibodies (**Fig. 2c**). Average counts from DSP profiling for the replicate 100 µm ROIs showed expression patterns that reflected the intensities seen with fluorescence microscopy for the corresponding target. To examine the other 41 oligo-conjugated antibodies in the cocktail, replicate 100 µm ROIs were plotted across cell lines with expected expression annotated as positive, negative, or ambiguous (**Fig. 2d, Supplementary Fig. 1**). In general, the counts obtained had a wide dynamic range, with negative and positive cell lines showing various levels of differential expression. CD20 and PD-L1 are examples of the two extremes in performance. The CD20 highest count average was 11,380 (SUDHL4) and the LOD in the negative cell lines (average counts plus 3 times the standard deviation) was 173.2, which is a 65.7-fold difference in counts between the highest expressor and the LOD. On the other hand, for PD-L1, the highest count average was 75.5 (SUDHL1) and the LOD was 30.8, which is only a 2.5-fold difference between the highest expressor and the LOD. These differences are likely reflective of differential abundance of these targets in these control cell lines and the intrinsic sensitivity and specificity of each antibody.

### ROI-to-ROI and section-to-section reproducibility

Since quantification of PC-oligos is performed using single-molecule counting, the digital counts for each analyte can be easily compared for digital precision. Unsupervised clustering of scaled data across all 5934 datapoints was performed to assess reproducibility across replicate ROIs from the cell pellet TMA (**Fig. 2e**). The six ROIs selected for each cell type clustered together across the entire dataset, verifying the reproducibility of different ROI sizes and replicates for this slide. The average R^2^ correlations of log transformed counts across the six ROIs for each cell line were determined by comparing the first replicate of the 100 µm ROI to the five additional ROIs. These five independent R^2^ values were averaged to obtain the values shown across the heatmap annotation (**Fig. 2e**). These average R^2^ values were all greater than 0.96, with a range from 0.963 (A549) to 0.999 (MOLT4). Expected clustering of similar cell line types was also seen with clustering of the leukemia and lymphoma cell lines (JURKAT, CCRF-CEM, HUT78, MOLT4, K562, SUDHL1, SUDHL4, DAUDI, RAJI) and the breast cancer cell lines (MDA-MB-468, SKBR3). Targets also tended to cluster by protein target type. For example, clustering was observed for the housekeeping proteins: Histone H3 and Ribosomal Protein S6 (S6), the T cell markers: CD3, CD4, and CD8, and the negative control IgGs. PD1, a target that is not expressed in any of the analyzed cell lines, also clustered with the negative control IgGs, showing antibodies with no specificity resemble the behavior of antibodies on samples lacking the expression of target epitopes. We also examined section-to-section reproducibility of 25 co-registered ROIs across two serial tonsil sections (**Fig. 2f**). Comparison of all antibodies across all ROIs had an of R^2^=0.97 and CD3, CD20 and PanCK antibodies were all greater than R^2^=0.93 (**Supplementary Fig. 2**), further reflecting the quantitative digital nature of the DSP methodology in tissue samples.

### Correlation of DSP counts with titrated cells

We next generated a cell pellet TMA to test the precision of DSP to quantify varying amounts of cells within an ROI. For this, two cell lines with differential expression of CD3 were mixed per core. CCRF-CEM cells (CD3-positive) were titrated into a background of HEK293 cells (CD3-negative) at the approximate percentages of CCRF-CEM cells at 100%, 95%, 70%, 30%, 5%, 0% (**Fig. 2g**). Four replicate 100 µm circles were analyzed across cell pellet titrations (**Fig. 2h**). Counts were plotted against the number of CD3-positive CCRF-CEM cells across the different cell pellet titrations ROIs. Targets expressed exclusively in CCRF-CEM showed the expected linear decrease in counts with the decrease in CCRF-CEM cells profiled. CD3, CD45, and CD45RO had R^2^ values of 0.95, 0.93, and 0.80, respectively. Importantly, targets expressed in both cell lines, RPS6 and AKT, did not show a similar decrease in counts with R^2^ values of 0.40 and 0.01, respectively. Mouse IgG2a also showed no trend, with relatively low counts across all ROIs. This data demonstrates the capability of the DSP platform to quantify varying numbers of cells across ROIs.

Since this mixed-proportion cell pellet assay revealed strong correlations between observed counts above background and positive cell numbers in an ROI, this enabled us to calculate LODs for targets expressed exclusively in CCRF-CEM cells. The LOD of each target was calculated, and the intercept of this LOD with the linear regression trend line was calculated to estimate the numbers of cells necessary to detect each target (**Fig. 2h**). CD3, CD45, and CD45RO had LODs approaching single cell detection at 1.3, 2.0, and 3.7 cells, respectively.

### *In situ* detection of mRNA with DSP

We also developed an assay that enables quantitative *in situ* detection of RNA targets. The DSP RNA detection assay uses oligonucleotide probes containing sequences complementary to targets of interest and an indexing PC-oligo sequence (**Fig. 3a**). Numerous probes across the transcript, with identical indexing PC-oligo sequences, were used to increase the sensitivity and isoform coverage of the assay (referred to as “tiles” from here on). For the NGS readout, each individual tile contains a unique sequencing index sequence and is therefore independently counted.

**Fig. 3:**
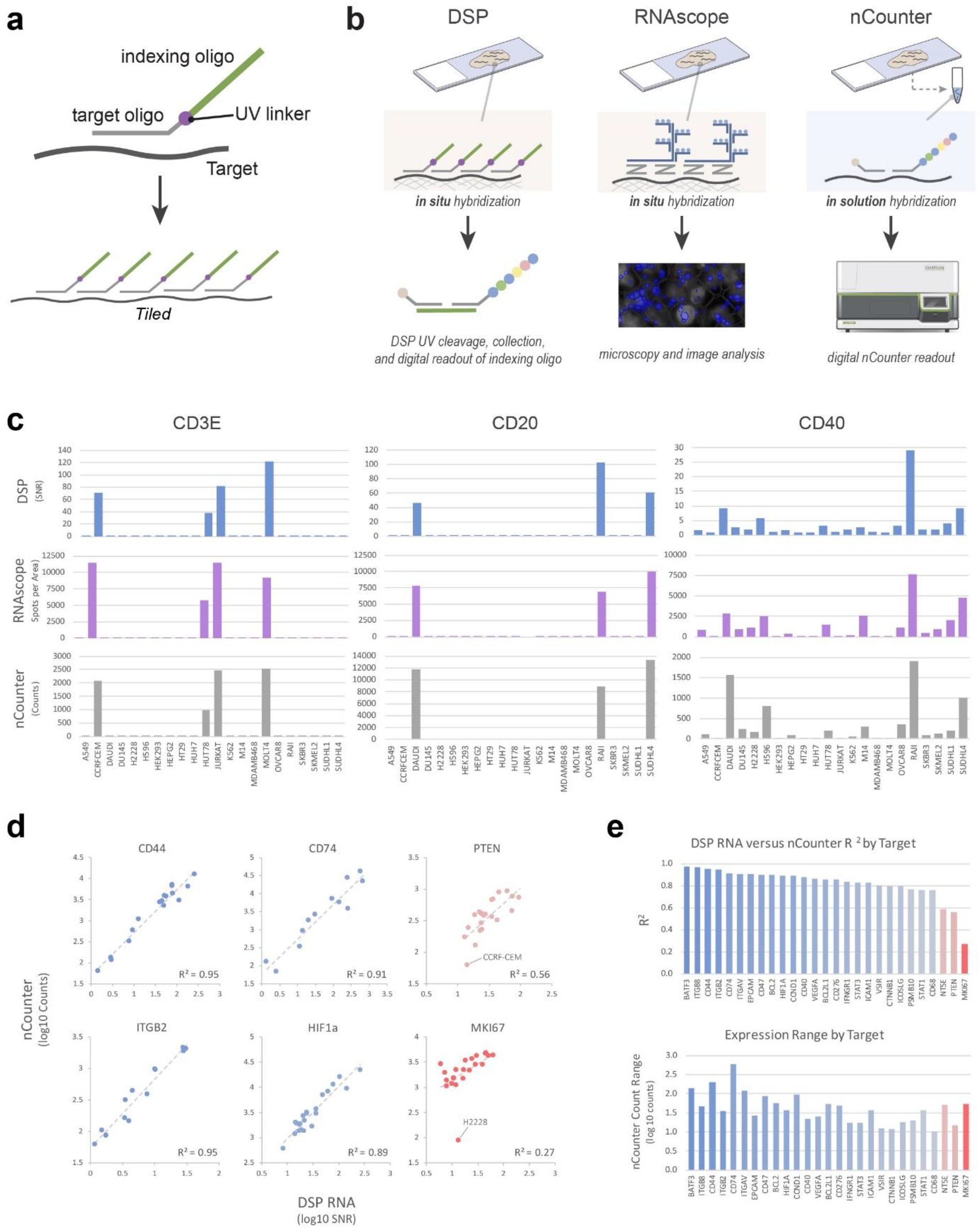
Validation of oligo-conjugated RNA probes for DSP. **a**, Oligonucleotide probes were designed with a target-specific complementary sequence joined to an indexing sequence by a UV-cleavable linker. To amplify signal, multiple RNA probes were designed to target various regions of each transcript (tiled). **b**, Overview of orthogonal approaches used for validation of RNA DSP. DSP, left: RNA targets (dark gray) are bound to tissue sections via formalin cross-linking. DSP probes bind directly to cross-linked targets (*in situ* hybridization) and the indexing portion of the probe is digitally counted after UV-cleavage. RNAscope, middle: In situ hybridization is performed with a series of amplifying oligos and a final layer allowing visualization with fluorescent probes (either directly conjugated to oligos, or through a tyramide signal amplification (TSA) reaction). Fluorescent spots, each representing individual transcripts, are quantified with image analysis software. nCounter, right: RNA transcripts are purified, hybridized in solution with nCounter barcodes, and counted on the nCounter platform. Note nCounter barcodes hybridize directly to RNA molecules, while for DSP, nCounter barcodes hybridize to the indexing PC-oligos. **c**, Comparison of CD3E, CD20, and CD40 transcripts across 22 cell pellet types with all three platforms. Top row: DSP SNR (target counts / average negative control counts) for an average of two 400 µm diameter circle ROIs (0.1320 mm^2^ total area). Middle row: RNAscope spot counts for two 300 × 400 µm ROIs (0.24 mm^2^ total area). Bottom row: nCounter PanCancer IO360 counts from 100 µg purified RNA from FFPE sections. **d**, Correlation of log transformed cell pellet data from DSP RNA analysis and nCounter data from purified RNA. Representative targets shown with at least 10 data points significantly above background for nCounter and covering at least one order of magnitude of expression. R^2^ values shown. Samples in blue have R^2^ >0.75 and samples in red have R^2^ ≤ 0.75. **e**, Summary of R^2^ values for all targets meeting criteria in d and the range of expression (highest nCounter counts / lowest nCounter counts across all cell pellets). Samples in blue have R^2^ >0.75 and samples in red have R^2^ ≤ 0.75.

### Cross platform validation on target subset

Since the accuracy of directly measuring RNA probe expression with primary or secondary detection approaches can be inaccurate and complicated on FFPE tissue, we tested the specificity of the DSP RNA assay against two orthogonal approaches, fluorescent *in situ* hybridization (FISH) and the nCounter System (**Fig. 3b**). FISH, direct detection of RNA transcripts on mounted FFPE sections, was performed using a commercially available RNAscope assay (ACD). Fluorescent detection of RNA transcript abundance from fluorescence microscopy images was quantified with QuPath image analysis software (**Supplementary Fig. 3**). A second approach for comparison was the use of direct detection and digital counting of whole slide purified RNA transcripts using the PanCancer IO 360 assay (IO360) on the nCounter platform (NanoString Technologies). The IO360 panel measures 770 transcripts, covering the majority of targets in the 88 immuno-oncology focused DSP RNA panel (**Supplementary Table 2**). The DSP RNA panel contains a total of 96 targets that includes targets for 8 non-specific RNA sequences, from a set of control sequences developed by the External RNA Controls Consortium (ERCC), to measure the background of the assay. ROIs of each unique cell pellet core of the TMAs were analyzed with DSP in duplicate. Digital counts of PC-oligos were counted using the nCounter platform. We observed highly concordant expression patterns with all three methods (**Fig. 3c**). In particular, CD3E, CD20 (MS4A1), and CD40 showed similar profiles across 22 human cell lines (**Fig. 3c**).

Further validation focused on the comparison of DSP RNA expression analysis versus the non-spatially resolved nCounter System readout of purified transcripts from whole tissue slide sections. For this, we focused our analysis on transcripts that were well above background in the nCounter System (at least 50 digital counts). Correlation of transcripts that were above background in 10 or more cells lines, with a dynamic range of expression >10-fold, were analyzed (**Fig. 3d**). The majority of targets analyzed (24 of 27) had R^2^ values above 0.75, with an apparently even spread of correlations values from 0.76 to 0.97 (**Fig. 3e**). CD44, CD74, ITGB2, and HIF1a scatterplots show examples of targets with high correlation from 0.89 to 0.95 (**Fig. 3d**). PTEN and MKI67 were the greatest outliers with each target having relatively low expression in a single cell line. This discrepancy could be explained by the known range of transcript variants for these targets, and the difference in probes used for the two assays, with the nCounter System readout only measuring the abundance of a single 100 nucleotide (nt) region, where the DSP assay readout is an aggregate of counts for 10 different ~35nt regions distributed across the transcript. Another explanation for this result is that PTEN and MKI67, markers for the cell cycle and proliferation, vary in expression depending on the density and number of cell divisions upon cell collection and RNA isolation.

### High multiplexing capabilities of RNA using NGS

We sought to increase DSP multiplexing capabilities of RNA by using next-generation sequencing as an alternative readout to the nCounter System. The NGS indexing oligo was redesigned to contain three features: (1) PCR primer binding sites for the addition of Illumina adapters and dual-indexing sequences, (2) unique molecular identifier (UMI) for the post-analysis removal of PCR duplicates, and (3) barcoded sequence for the identification of the RNA target. We generated a 928-plex NGS readout panel identical to the nCounter RNA panel except for the readout portion of the oligo. In contrast to the nCounter System indexing oligo, the NGS design allowed for the ability to resolve each tile along the RNA transcript (**Fig. 4a**). DSP analysis was performed on the 23-core cell pellet array using the nCounter or NGS panel (**Fig. 4b**). We found high cross-platform concordance (R^2^ > 0.8) among targets that show positive expression with the nCounter System (**Supplementary Fig. 4**), validating the NGS readout workflow. In particular, this demonstrated the accuracy of UMI-based digital counting strategy as compared to single-molecule counting by nCounter. NGS readout negative controls (glass surface only and mouse 3T3 cell pellet) showed significantly lower background counts compared to the 22 human cell lines (**Fig. 4c**). Furthermore, expression profiling of the 928-plex showed differential expression patterns between the cell pellets as expected (**Fig. 4c)**. For example, the T cell marker CD3E showed significantly higher expression in T cell derived cell lines (CCRF-CEM, HUT78, JURKAT, MOLT4) compared to the other 19 cell lines (**Fig. 4d**). A key feature of NGS readout compared to nCounter System readout is its ability to assess several independent measurements per RNA transcript target through the tiled probes. Interestingly, we observed cases where counts targeting specific exons were differentially expressed between cell pellets (**Fig. 4e**). These data suggest that DSP RNA with NGS readout has high-multiplexing capabilities as well as the potential to probe fundamental aspects of RNA biology including alternative splicing.

**Fig. 4:**
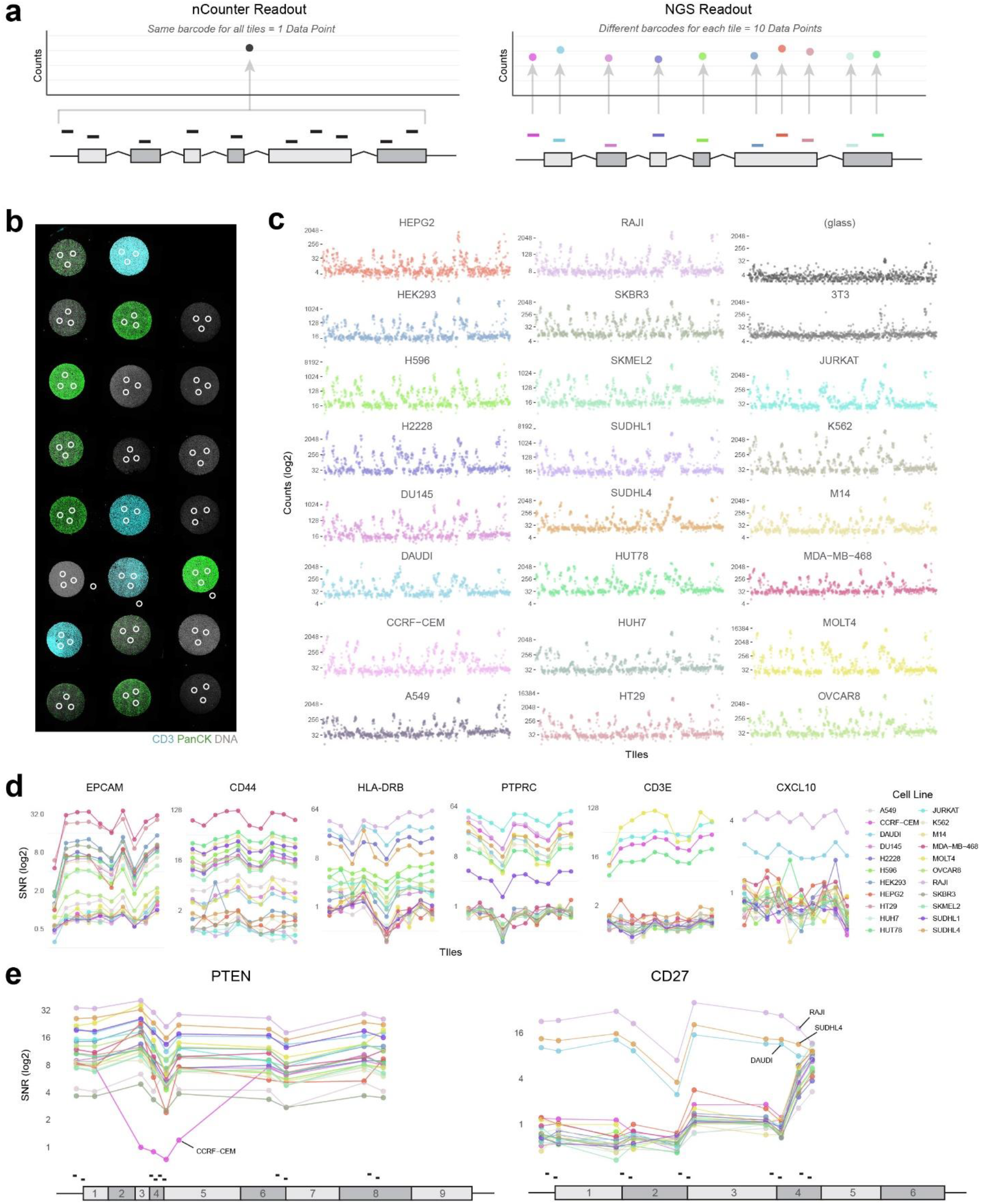
NGS readout allows high-plex analyses of each transcript. **a,** Comparison of nCounter and NGS readouts. For the nCounter readout, all tiles of a transcript are readout with the same barcode, resulting in one data point per target. For the NGS readout, each tile has its own unique barcode, resulting in a data point for each tile. **b,** ROI selection on the 23-core cell pellet array for NGS readout. Three 300 µm diameter circles were selected as replicates from each cell pellet. Three additional ROIs on the glass surface only were included as a negative control. **c,** Graphs of NGS readout counts from multiplexing 928 RNA in situ hybridization probes onto the 23-core cell pellet array. Each graph position corresponds to the cell pellet position in b. Glass negative control graph placed on the top right. Y-axis indicates NGS digital counts, with each point representing the average of the three ROIs replicates for each cell pellet. X-axis indicates each tile grouped by gene. For example, first 10 points represent 10 tiled probes along AKT1 transcript and cluster together in counts. **d,** Expression of representative transcripts across all cell lines. SNR (y-axis) calculated for each tile by dividing the average deduplicated NGS counts by the geomean of the negative probe counts in each sample. For each target, data from 10 tiles is shown 5’ to 3’, from left to right. Each point indicates an independent tile (x-axis) measurement for the transcript. For visualization purpose, lines were drawn between each independent point to distinguish between each cell pellet. **e,** Expression of two targets with differential expression across transcripts. Tile counts with corresponding exon position for PTEN (left) and CD27 (right). Line graphs of SNR with each point aligned above the tile position shown in the gene structure below the graph. For visualization purposes, lines were drawn between each independent point to distinguish between each cell pellet.

### Geometric profiling of protein and RNA

Our first thorough examination of this platform on FFPE tissue sections was performed with simple geometric profiling of lymphatic tissues, which have known expression of many targets in our IO protein and RNA panels, as well as a sampling of tumor tissues with unknown expression profiles. Profiling was performed on tonsil tissue by imaging 4-color fluorescence of T cells (CD3), B cells (CD20), Epithelial cells (PanCK), and DNA (SYTO 83) to distinguish tissue morphology (**Fig. 5a**). Twelve 200 µm diameter circular ROIs were selected based on enrichment of CD3, CD20, or PanCK for photocleavage of the 44-plex oligo-conjugated antibody cocktail (**Fig. 5a**, **Supplementary Table 1**). nCounter System counts for the 3 markers used to visualize ROIs, CD3, CD20 and Pan-CK, strongly correlated with each morphological enriched region (**Fig. 5b).** Unsupervised clustering of regions with similar visual profiles (enriched for each morphology marker) grouped together demonstrating the correspondence between local biological identity, such as germinal centers and the high-plex molecular profiles based on B cells, T cells and epithelium. A similar approach was used to profile colorectal cancer tissue. Circles with 665 µm diameter were selected based on localization at the invading margin, center of the tumor or in adjacent normal tissue was performed by imaging 4 immunofluorescent morphology markers including T cells (CD3), total immune cells (CD45), tumor and normal epithelial cells (PanCK), and DNA (SYTO13), as well as the cocktail of 44 PC-oligo conjugated antibodies (**Fig. 5c**). Unsupervised clustering of the nCounter System counts from these geometric ROIs were examined (**Fig. 5d**) showing that ROIs clustered according to spatial immune biology of the tumor at the invading margin, the center of the tumor, and normal tissue. As would be expected, the invasive margin expressed the highest levels of immune cell-related markers including those associated with an adaptive (T cells, B cells) and innate (myeloid) immune response, while the tumor center was associated with tumor markers such as PanCK, beta-2-Microglobulin (B2M) and proliferation (Ki67).

**Fig. 5:**
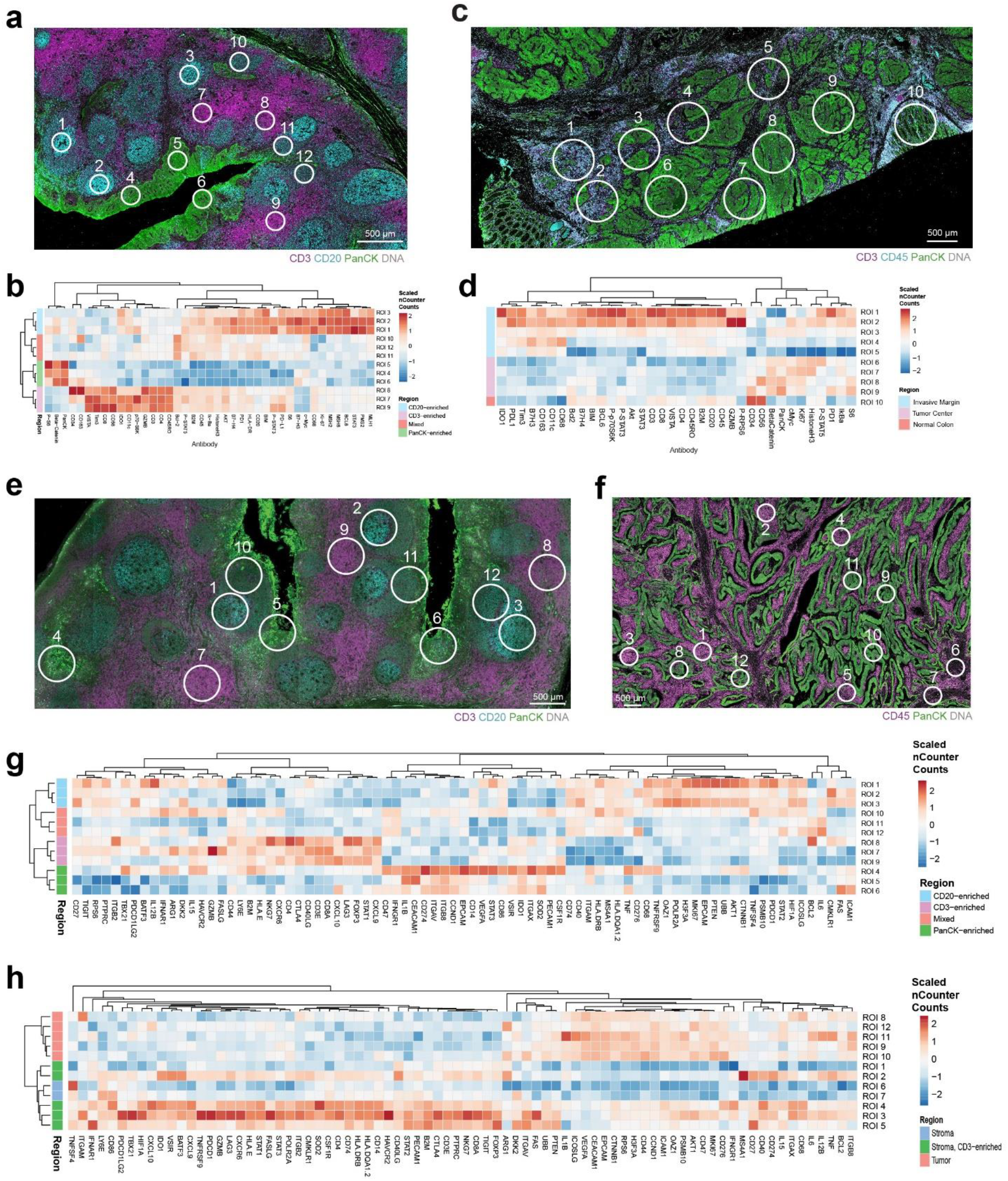
Spatially-resolved, multiplexed protein characterization of geometric regions on FFPE tissue. **a**, Protein profiling of tonsil: A tonsil section was imaged using 4-color fluorescence of CD3 (magenta), CD20 (cyan), PanCK (green), and DNA (grey) to establish overall tissue morphology. Twelve 200 µm diameter circular ROIs were selected for detailed molecular profiling with a 44-plex oligo-antibody cocktail. **b**, Unsupervised hierarchically clustered heatmap of region-specific nCounter counts across all protein targets on tonsil. Regions with similar visual profiles cluster together, demonstrating the correspondence between local biological identity and molecular profiles. Region annotations were manually curated based on immunofluorescence staining patterns. **c**, Protein profiling of colorectal cancer (CRC): A CRC section was imaged using 4-color fluorescence of CD3 (magenta), CD45 (cyan), PanCK (green), and DNA (grey) to establish overall tissue morphology. Ten 665 µm diameter circular ROIs were then selected for detailed molecular profiling with a 44-plex oligo-antibody cocktail. **d**, Unsupervised hierarchically clustered heatmap of region-specific nCounter digital counts across all protein targets on CRC tissue. **e**, RNA profiling of tonsil: A tonsil section was imaged using 4-color fluorescence of CD3 (magenta), CD20 (cyan), Pan-CK (green), and DNA (grey) to establish overall tissue morphology. Twelve 300 µm diameter circular ROIs were selected for detailed molecular profiling with a 96-plex RNA probe cocktail. **f**, RNA profiling of CRC: A CRC section was imaged using 3 color fluorescence of CD45 (magenta), PanCK (green), and DNA (grey) to establish overall tissue morphology. Twelve 300 µm diameter circular ROIs were then selected for detailed molecular profiling with a 96-plex RNA probe cocktail. **g**, Unsupervised hierarchically clustered heatmap of region-specific nCounter counts across all RNA targets on tonsil that had at least one ROI above background. Regional annotations based on fluorescence are labeled as CD3-enriched, CD20-enriched, mixed and PanCK-enriched. **h**, Unsupervised hierarchically clustered heatmap of region-specific nCounter digital data across all RNA targets on CRC that had at least one ROI above background. Regional annotations based on fluorescence are labeled as stroma-enriched, CD3-stroma-enriched and PanCK-enriched.

We also performed geometric profiling of *in situ* RNA expression in tonsil and colorectal cancer tissue (**Fig. 5e, 5f**). Tissue-specific morphological features were highlighted with fluorescently-labeled antibodies following *in situ* hybridization, PanCK and DNA (SYTO13) were used for all tissues and CD3, CD20 or CD45 were used for tonsil and colon tumor tissue, respectively (**Fig. 5e, 5f**). All samples were hybridized with the 96-target human IO RNA panel (**Supplementary Table 2**). In the tonsil tissue, 300 µm diameter circular ROIs were selected to include epithelium, T cells, B cells or mixed populations, similar to the ROI selection strategy performed for protein. ROIs from colorectal cancer samples were selected in the tumor or stroma. Unsupervised hierarchical clustering of all tonsil ROIs across all targets showed that T cells, B cells and epithelial regions grouped together based on local biology for each cell type (**Fig. 5g**). ROI selection for the colorectal cancer tissue was done slightly different than for protein, where regions were selected based on tumor-, stroma-or stroma and CD3-enriched areas. Unsupervised clustering nCounter counts of colorectal cancer ROIs showed a strong separation of tumor and stromal ROIs, with the stromal CD3-enriched ROIs being intermixed with the total stroma (**Fig. 5h**).

Finally, we demonstrated the high throughput capability of DSP by profiling 44-protein targets across a 384-core (665 µm diameter circles per core) colorectal tumor tissue microarray (TMA) (**Supplementary Fig. 5**). This high number of geometric cores was collected on the DSP instrument in one day. We found that targets profiling similar cell types had the best correlation across all cores, as would be expected. Furthermore, two replicate cores from one patient (of 192 patients’ cores) had high levels of PD-L1 relative to all other samples, highlighting the capability of DSP to detect rare events with high throughput sample characterization.

### Gridded profiling of protein and RNA

We used the DSP instrument to profile tissue in a “gridded” manner, with square ROIs arrayed directly adjacent to each other for deep expression profiling of one concentrated area. This gridded-profiling was performed on the same tonsil sample used for geometric protein profiling (**Fig. 5a**) also with 4-color fluorescence of CD3, CD20, PanCK, and DNA. A total of 196 gridded ROIs of 100 µm × 100 µm size (~170 cells per ROI) were selected (**Fig. 6a**). As would be expected, the distribution of each morphology marker could directly be recapitulated by spatially distributing nCounter counts for CD3, CD20 and Pan-CK, respectively (**Fig. 6b**). The additional 44 PC-oligo antibodies also showed distinct spatial patterns of protein expression with different morphology visualization markers (**Fig. 6c and Supplementary Table 1**). Morphology-related patterns were observed for CD4, CD8, CD3, B7-H3, PD-1, and Ki67 counts, while HLA-DR had similar spatial patterns to CD20. Beta-catenin had similar spatial profiles as PanCK, as both are related to epithelium. VISTA and Bcl2 had a unique spatial profile distinct from the three morphology visualization markers. A majority of the 44 PC-oligo conjugated antibodies showed distinct spatial profiles, except two housekeeping proteins, Histone H3 and S6, which should be expressed relatively equally according to the number of cells and isotype controls (mouse and rabbit IgG), which were evenly distributed across the grid, giving a “static-like” profile, which was not related to biology within the tonsil. (**Supplementary Fig. 6**).

**Fig. 6:**
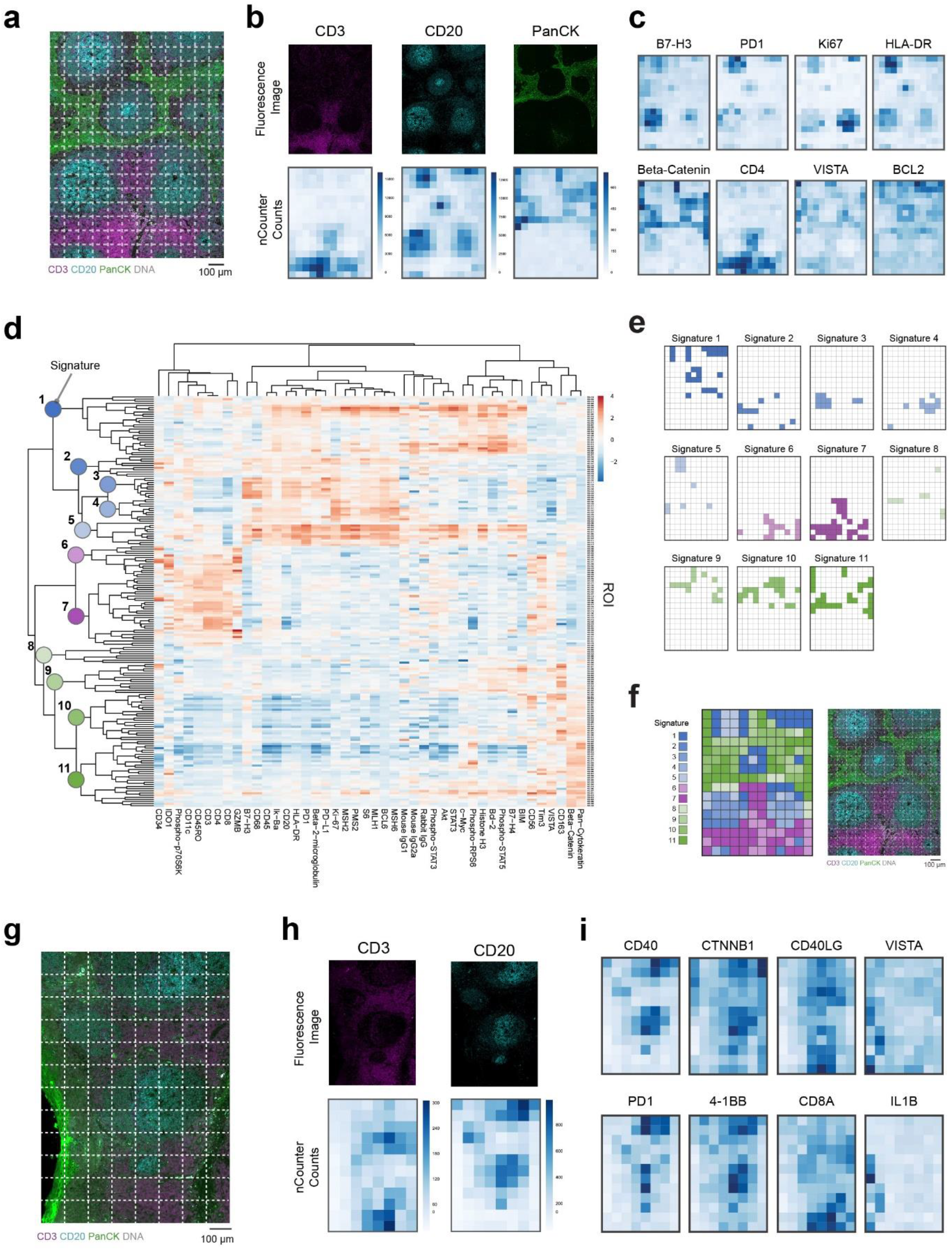
Protein and RNA profiling with gridded ROIs. **a**, Protein profiling of tonsil: A different section of the same tonsil from Fig. 4 was imaged using fluorescence of CD3 (magenta), CD20 (cyan), PanCK (green), and DNA (grey). Gridded profiling was performed by selecting 192 adjacent 100×100 µm square ROIs for detailed molecular profiling with a 44-plex oligo-antibody cocktail. **b**, Heatmap of nCounter counts obtained for CD3 (magenta), CD20 (cyan), Pan-CK (green) for all ROIs compared to single channel visualization images for the same target, respectively. **c**, Heatmaps for protein counts for other targets show distinct patterns of expression across the grid, which are similar to the fluorescent visualization markers. Entire dataset for this one ROI contains spatial information for 44 different antibodies, all with unique expression profiles. **d**, Unsupervised hierarchically clustered heatmap of all ROIs and targets. Specific nodes highlighted as unique signatures that were further analyzed. **e**, ROIs within each signature highlighted in a grid format matching the grid used for DSP profiling. **f**, All signatures combined, next to original fluorescent image for comparison **g**, RNA profiling of tonsil: A different section of the same tonsil from Figure 4 was imaged using fluorescence of CD3 (magenta), CD20 (cyan), PanCK (green), and DNA (grey). Gridded profiling was performed by selecting 96 adjacent 100×100 µm square ROIs were then selected for detailed molecular profiling with a 96-plex RNA probe cocktail. **h**, Heatmap of digital counts obtained for CD20 and CD3 across ROIs compared to single channel images of visual staining markers for the same target. **i**, Heatmaps of RNA counts for other targets show distinct patterns of expression across the grid, which are similar to the fluorescent visualization markers.

For an unbiased analysis of this gridded data, hierarchically clustered heatmap analysis was performed across all ROIs and targets (**Fig. 6d)**. Specific clusters, or nodes, were selected and highlighted as unique “signatures”. ROIs within each signature highlighted in a grid format matching the grid used for DSP profiling (**Fig. 6e)** and combined (**Fig. 6f)**, displaying tissue structures remarkably similar to those seen in the original four-channel immunofluorescence image.

Analysis was also performed on tonsil tissue using the *in situ* IO RNA panel and similar results where RNA expression counts mimicked morphology patterns based on distinct immune biology within the tonsil (**Fig. 6. g, h, i**).

### Segment profiling of tumor and surrounding microenvironment

To demonstrate the capability of DSP to capture biology based on the contexture of the organic tissue structure, we further characterized the colorectal tumor geometrically profiled in **Fig. 5c** by segmented profiling to directly assess protein expression in two separate compartments of the TME, tumor cells, and the surrounding microenvironment, such as stroma and immune cells (**Fig. 7a**). To assess segment profiling, it was essential to visualize tumor cells by PanCK expression. Based on the PanCK-positive tumor within the same geometric 665 µm diameter ROIs is shown in **Fig. 5c**, PanCK-expressing regions with the ROI were exposed to UV and collected, followed by UV illumination of the tumor inverse area, or PanCK-negative, within the same ROIs (**Fig. 7a)**. Selection of PanCK-positive cells can be done automatically within the DSP using the DSP ROI selection software (**Fig. 7b** and *data not shown*). A heatmap of tumor (PanCK-positive) and tumor inverse (PanCK-negative) segments (**Fig. 7c**) revealed that two distinct expression profiles, based on each segment, with a strong enrichment of immune markers in the tumor inverse or stromal compartment. The tumor compartment of ROI1, which is located at a highly immune-infiltrated region of the invasive margin, showed the greatest enrichment of immune markers, compared to other tumor compartments, in the tumor center, but also other ROIs in the invasive margin. This increased expression of immune markers is likely due to high prevalence of immune infiltrates within this region of the tumor. ROI overlayed heatmaps of tumor inverse segments is shown to demonstrate that high, but varied expression of CD3 and CD11c are observed in the surrounding microenvironment at the invasive margin, but CD34 is down-regulated in those same regions (**Fig. 7d**).

**Fig. 7:**
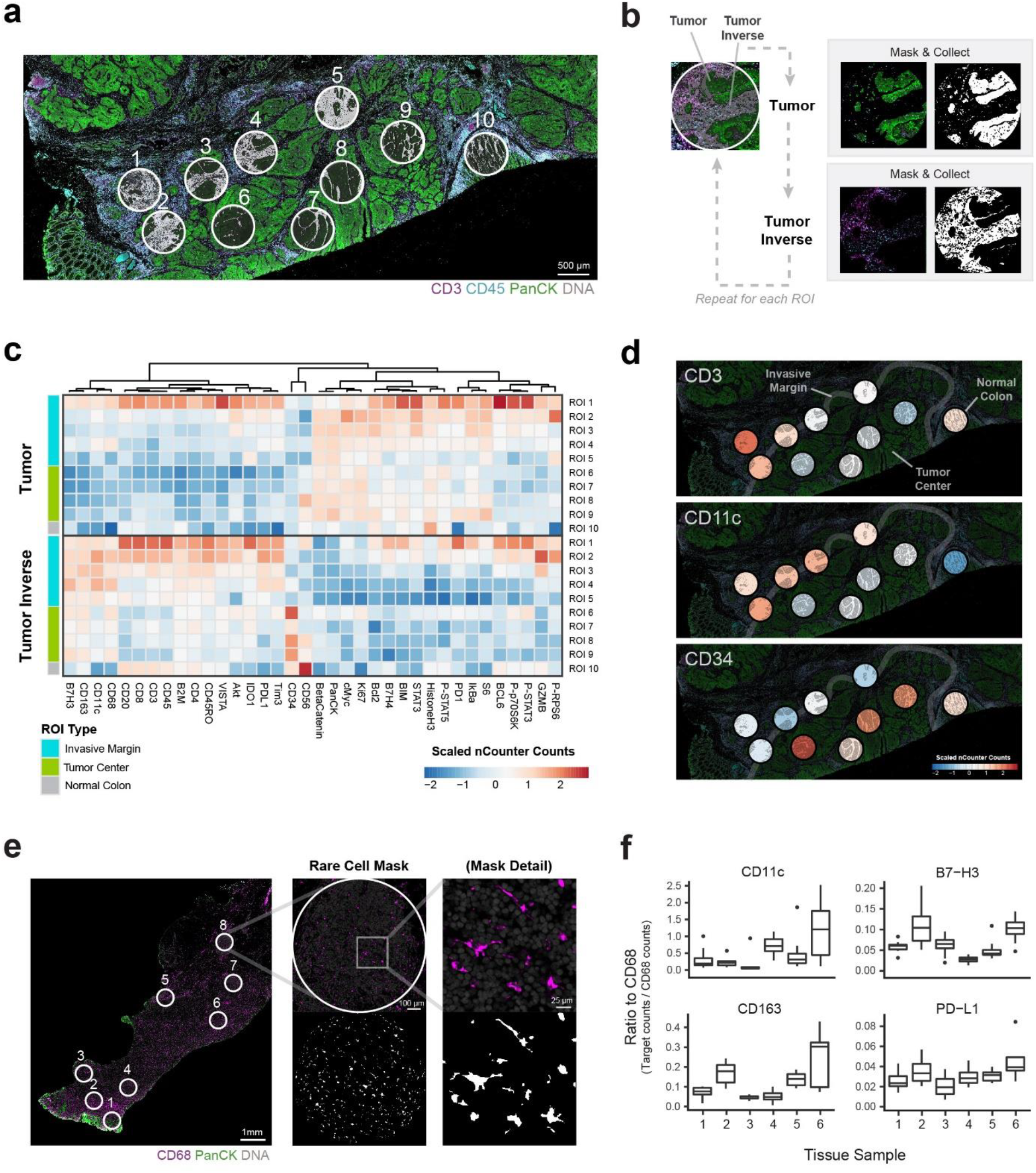
Segmented and rare cell profiling of different compartments of colorectal tumor and IBD tissues. **a,** Segmented profiling on the same ROIs on CRC shown in Fig. 5b. Two segments were profiled within each 665 µm diameter circular ROI: Pan-CK positive area shown in black (the tumor, or normal epithelial) and the remaining Pan-CK negative area of the ROI (inverse). **b,** Workflow for collection of multiple segments within the same ROI. Segments are first identified (tumor and tumor inverse). The PanCK fluorescent images is used to create a mask for only Pan-CK positive cells, UV is illuminated within only this mask, and the eluate is collected and dispensed in a microtiter plate. The slide is washed, to remove any residual PC-oligos, and the procedure is repeated for the inverse of the Pan-CK mask. This workflow is then repeated for each ROI across the sample. **c,** Hierarchical clustering of ROIs arranged by location or type (invasive margin, tumor center, or normal colon) and by segment type (tumor or tumor inverse). **d,** Heatmap of tumor inverse segments for selected targets. Landmarks are annotated to highlight the relationship between target counts and the location of each ROI. **e**, Rare cell profiling of IBD tissue: Example of eight ROIs and masking scheme used to profile CD68-positive (macrophage) segments. **f**, Six IBD samples profiled as described in e. nCounter counts for all targets were normalized to CD68 counts. Differential expression of macrophage biological markers: CD163, B7H3, CD11c and PD-L1 vary between different IBD samples.

### Rare cell profiling and contour profiling inflammatory bowel disease

Since autoimmune diseases that manifest in the gastrointestinal system, such as inflammatory bowel disease (IBD), have diverse pro-inflammatory immune cells, we used DSP to characterize rare cells and areas adjacent to immune cells in the IBD tissue microenvironment. We stained samples for macrophage (CD68), T cell (CD3), epithelium (PanCK) and DNA (SYTO 83) and the 44-PC-oligo-conjugated antibody cocktail to interrogate IBD regions containing concentrated macrophage populations (**Fig. 7e, f, Fig. 8**). We used two approaches for assessing macrophage biology in IBD samples: 1) Unique cell type profiling of CD68 cells in various inflamed regions of the tissue (**Fig. 7e, f**), and 2) Contour profiling of markers at varying distances from macrophage-enriched regions (**Fig. 8**).

**Fig. 8:**
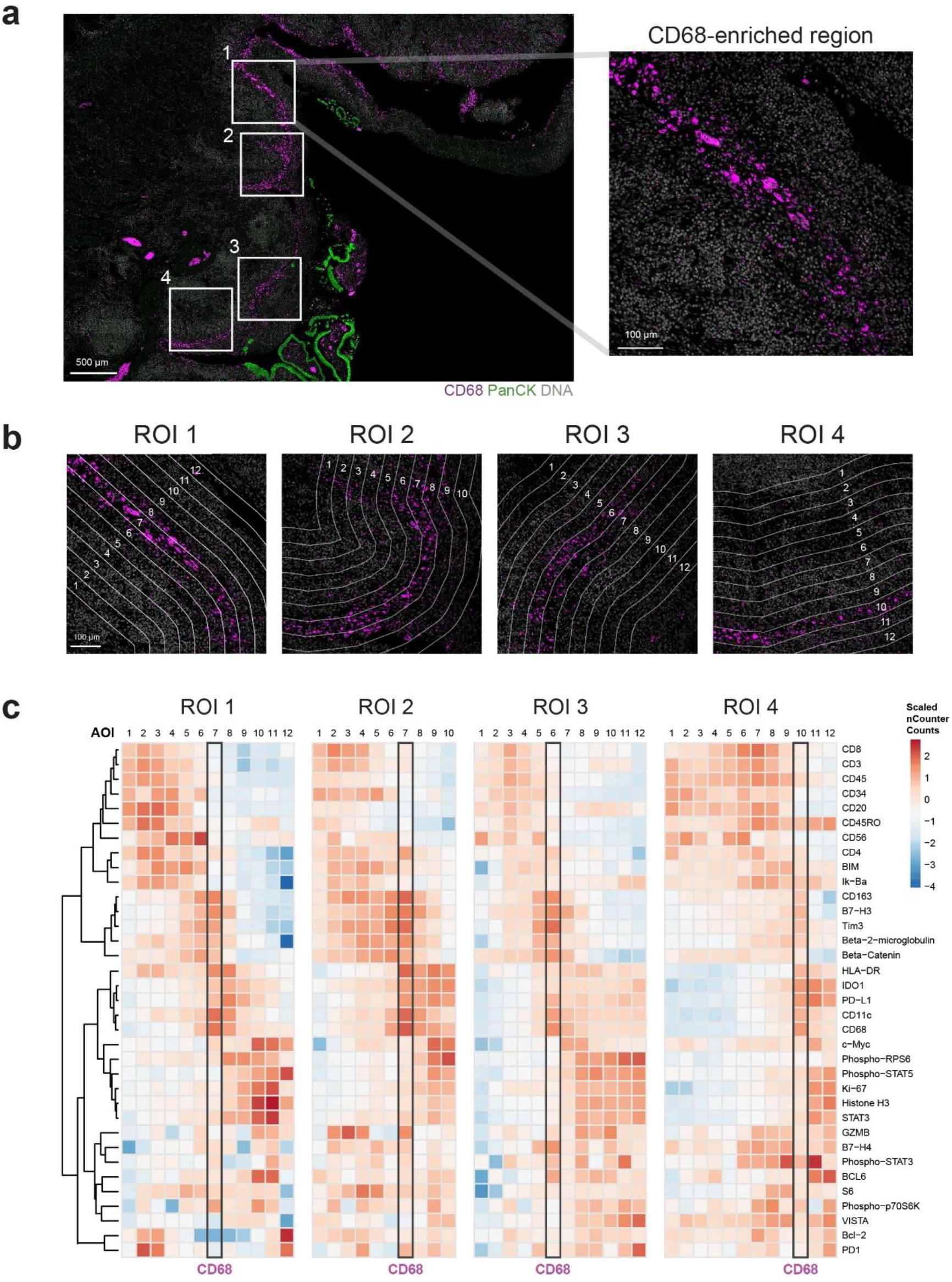
Contour profiling of a macrophage-enriched region of inflammatory bowel disease. **a**, An IBD sample was used to select four 665 µm × 665 µm square ROIs that contained a CD68-enriched trail of cells. **b**, Within each ROI contour selection based on the CD68-enriched trail of cells was segmented by a 50 µm region. From each CD68-enriched area, additional segments with widths of 50 µm were selected from each side toward the lumen or epithelium. Each segment was collected and analyzed for 44-plex protein markers. **c**, Unsupervised hierarchical clustering markers in each ROI. Black boxes show the CD68-enriched region.

For rare cell profiling, 6 different IBD samples were processed as described above and macrophages were identified using fluorescently labeled CD68 antibody, to be illuminated with the DMD on the DSP system. This rare cell profiling of macrophages in IBD showed that some samples had higher wound healing pro-inflammatory macrophage markers (CD163, B7-H3), while other samples had antigen presenting profiles (CD11c) and other markers had no enrichment (PD-L1). This showed that different samples and regions had unique macrophage biology that was only captured by multiplexing immune markers.

One of the IBD samples from the 6 tissues used for rare cell profiling displayed an interesting pattern of macrophages (CD68) that followed the tissue architecture of the colorectal epithelium. To further assess the biology of this sample, we also performed contour profiling to measure protein markers that were expressed near the macrophage-enriched areas. The slide was processed with fluorescent morphology markers for macrophages/CD68, epithelium/panCK and DNA/SYTO 83, as well as the 44-oligo-conjugated antibody cocktail. 665 µm × 665 µm ROIs were selected that contained ribbons of macrophages that are coordinated with the colon epithelium (**Fig. 8a**) and assessed biology related to macrophage using contour profiling of markers at varying distances from the macrophage-dense region. Within each ROI, the macrophage segment was contour plotted according to the CD68 morphology marker and additional segments, each 50 µm distance from the CD68-dense region (**Fig. 8b**). Hierarchical clustering of digital counts normalized to ROI area for each marker (**Fig. 8c**) clearly distinguished three clusters of immune biology based on proximity to the macrophage enriched region. For all samples: 1) moving toward the lumen (left side of the CD68 region) there were higher amounts of T cells (CD8, CD3, CD4, CD45RO) and NK cells (CD56) effector cells, 2) as expected, the regions closest to the CD68 dense segments expressed a number of macrophage (CD163, CD11c) and myeloid cell markers and activation/inhibitory myeloid markers (HLA-DR, B7-H3, PD-L1, IDO1), and 3) moving closer to the epithelium (right of the CD68 region) there was high expression of activation/effector molecules (PD1, VISTA, GZMB), proliferation (Ki67) and STAT signaling (pSTAT3, pSTAT5). Using this approach, we were able to identify three distinct clusters of immune biology based on proximity to a CD68/macrophage dense area. Large amounts of inflammatory T cells in the lumen of a colon are likely the cause of IBD, while myeloid cells act as a barrier and may be presenting self-antigens that activate T cells to cause IBD. Lastly there is clearly effector and cytokine related signaling in the epithelium of the IBD samples that may also be part of the dysregulated inflammatory response.

## DISCUSSION

Methods for multiplexed spatial analysis of FFPE samples have lagged far behind modern genomic high-plex techniques for molecular profiling being limited in the number of markers that can be simultaneously analyzed (typically under 10-plex). In this study, we demonstrate a new methodology, DSP, that combines modern genomic detection technologies (NGS and Digital Optical Barcodes), organic chemistry (photocleavable oligonucleotides), semiconductor technology (dynamically configurable digital micromirror chips), and microfluidic sampling to advance classic IHC to a modern high-plex genomic and proteomic profiling capability. We show that DSP can accurately detect over 40 proteins and over 900 mRNA probes. The NGS readout mode also has unlimited multiplexing ability and, it should be emphasized, that no changes in the hardware utilized on this microscope need to be altered to measure “unlimited” targets in a single FFPE tissue slice.

Since DSP uses automated microfluidics (microcapillary capture of PC-oligos combined with automated digital detection of the PC-oligos), it allows for an high dynamic range of expression (**Fig. 2e**)^13^. Protein and RNA was detected in a reproducible manner with high correlation to standard IHC and FISH, respectively (**Fig. 2f, Fig. 3**). The digital counts read by the DSP platform are also highly linear, and measurements of protein expression align well with fluorescent IHC images, while RNA expression correlates well with orthogonal expression profiling technologies (**Fig. 2c, 3c, 6b, 6h**). With the high quality of reproducibility and quantitation, advanced quantitative modes of genomic and proteomic molecular profiling should now be possible (*e.g*, establishment of quantitative biomarker profile cut-points, multi-site shared-data consortium studies, or all-digital spatially resolved reference databases).

A key component of the DSP instrument is the use of programmable DMD to perform the spatially resolved molecular profiling. This technology allows regions of any shape or size, including non-contiguous areas, to be analyzed down to single-cell resolution (**Fig. 7e**). The DSP can dynamically configure DMDs (~2 million total mirrors) for molecular profiling uniquely for each tissue sample and ROI to maximize the biological information obtained from that specific tissue slice. ROIs can be defined by regular or geometric boundaries (for example, a grid across a tissue microarray, **Fig. 5 and Supplemental Fig. 5**) or irregular boundaries (for example, regions defined by cell or tumor borders, or proximity to specific tissue structures, **Fig. 7, 8**). Notably, size, shape, or spatial distribution of the ROI does not alter sample collection time.

One major application that utilizes DMD morphology-based ROI selection is for studying immuno-regulatory proteins within the TME. The DSP platform allows automatic quantification of tumor (PanCK) and stroma (non-PanCK) in the TME using DMD specific fluorescent antibody guided selection of ROI selection (**Fig. 7a-d**). This allows for understanding the heterogeneity within the tumor and stromal compartments. Additionally, the DMD in the DSP platform allows for selection of hundreds of single-cells to be multiplexed in a single DSP read-out (**Fig. 7e-f**). The relatively low-plex fluorescence-based markers used in DSP provide full-tissue scan at single cell resolution that can be selected for collection in a single ROI. This allows for high-plex analysis of a single cell type population based on location within the tissue. For example, the tumor or IBD microenvironment can be evaluated in ROIs in key areas such as the tumor, immune infiltrate, stroma, and tumor-infiltrate boundaries (**Fig. 7a-d**).

A limitation of protein profiling technologies is that they generally depend upon FFPE-compatible antibodies, which can be difficult to generate for some targets with low expression or low immunogenicity. Additionally, as with other antibody-based competing assays, antibody efficacy can change dependent on antibody lot, age, tissue, type, and fixation parameters, which can lead to variation in digital counts between independent experiments. Normalizing digital counts to the same range across experimental slides can help control for antibody-dependent variation. Utilizing spatial mRNA profiling provides an alternative for cases where the necessary antibodies do not accurately detect antigen or exist at all. There are also important cases where the biology of the system generates divergent mRNA and protein spatial patterns, such as post transcriptional control by miRNA or post translational modifications that alter protein stability. Simultaneous RNA and protein detection could reveal novel mechanisms of regulation that could be targeted therapeutically. Transcript-specific probes can also be made rapidly, in a high-throughput manner, using synthetic oligonucleotides, allowing for fast, completely custom high-plex panels to be easily developed. DSP RNA readout chemistry using NGS, allow each individual tile to be quantified independently, enabling the investigation of genes with differential abundance of expression across the transcripts and also allow spatial resolution of heterogenous differentially spliced isoforms.

To facilitate the simultaneous analysis of spatially resolved RNA and protein, we designed the GeoMx DSP software to allow one to overlay the ROIs from one tissue section (performed using high plex protein panel) directly onto the spatially aligned adjacent tissue section utilized for high-plex RNA profiling (*data not shown*). The Proteinase K digestion step utilized for *in situ* RNA hybridization destroys some of the key epitopes for antibody binding rendering simultaneous protein and RNA on a single section viable only for relatively high-abundance protein targets (*e.g.,* Pan Cytokeratin). By aligning the adjacent sections images together via software, and then projecting the ROI geometries (from protein to RNA or vice versus), rigorous simultaneous analysis (*e.g.,* differential expression volcano-plots, principle component analysis) can be performed on both digital layers of information. This type of “projection-combination” of protein and RNA data works for all modes of detection except for rare-cell and single-cell modalities.

Some limitations of the DSP platform should be noted. Profiling every cell in a tissue slice at single-cell resolution with this system is impractical and cost-prohibitive. One advantage of imaging-based methods over DSP, is the ability to get multiplexed information on each cell in the tissue slice, whereas DSP provides high-plex information of regions-of-interest and tissue compartments-niches. Hence, we view DSP and imaging-based approaches to be complementary, and in-fact, cycling-based imaging approaches can be utilized up-front of a DSP workflow for those cases where high-plex guided region-of-interest selection for DSP is desired. DSP is, perhaps, most closely aligned with Laser Capture Microdissection methodologies, but DSP has the advantage of being high-throughput, totally non-destructive to the sample, and completely automatable.

DSP offers a robust, flexible platform to enable novel insights in spatially complex biological systems and to accelerate drug development. For example, experiments with tumor tissue sections and tumor microarrays reveal significant intra-and inter-tumoral heterogeneity of immune markers (**Fig. 7a-d and Supplemental Fig. 5**). Such data could be used to better understand how drug candidates modulate the tumor microenvironment and affect processes such as immune cell activation and recruitment. Fundamental questions about tumor development, heterogeneity, response to immunotherapy and factors affecting growth and metastasis can be explored. Beyond oncology, infectious disease researchers could assay heterogeneous structures such as granulomas to profile host-pathogen interactions. The technology also could benefit many fields, such as developmental biology and neurobiology, that require detailed characterization of the spatial distribution of cells in tissues.

The GeoMx DSP platform also has considerable application for translational research, particularly in the IO field, due to the extreme importance of increasing our understanding of tumor immune status ^22^ and attempting to predict clinical responses to immunotherapies. Indeed, while it is clear that IO agents have revolutionized cancer therapy, immunotherapies are also associated with significant toxicities (*e.g.* IBD). Biomarkers able to identify tumors from patients most likely to respond to a given therapy and distinguishing those that have adverse side effects still remain elusive. The DSP platform allows for automatic quantification of tumor and stroma in the TME (**Fig. 7a-d**), as well as, rare localized cell types within different locations in a tumor or toxicity sites (**Fig. 7e-f**). This will aid the understanding heterogeneity within the tumor and stromal compartments is increasingly necessary for tumor stratification and drug discovery^23, 24^ but remains a key challenge in the development of targeted cancer therapeutics^25^.

The simplicity of using the GeoMx DSP platform makes it easily optimized for future clinical applications. It can be easily run in any clinical pathology lab and outcompetes comparable technologies in most categories important for clinical use. DSP does not require any complex detection instrumentation (*e.g.,* mass spectrometer), is highly automated, and routinely operated by non-specialized personnel. Anyone familiar with performing simple tissue-based immunohistochemistry can operate the GeoMx System, with automation providing the user with step-by-step guidance. Furthermore, DSP is non-destructive, and precious clinical specimens can therefore be stored and re-probed at any time. Additionally, companion diagnostic development is greatly enhanced since the higher multiplexing capability of the platform allows a much more complete mechanism-of-drug-action study to be performed. DSP brings the power of classic mid-plex and high-plex molecular profiling to spatial profiling. Hence the multiplexed molecular signatures that have been so powerful in predicting response to therapy can now be examined in their spatial context, greatly increasing the ability to predict which patients are likely to benefit from specific types and combinations of immunotherapy.

## METHODS

### Microscope and fluidics system overview

For most of the data presented in this paper (**Fig. 2a-f, Fig.3-7**), an automated imaging and sample collection instrument was developed by modifying a standard microscope and controlling it with Metamorph (Molecular Devices) and custom software. For protein or RNA detection, a multiplexed cocktail of primary antibodies or RNA binding probes, each with UV photocleavable indexing oligos, and/or 1-4 fluorescent markers (antibodies and/or DNA dyes) was applied to a slide-mounted FFPE tissue section (staining protocols described in the following sections). Antibodies and RNA probes used in these studies are listed in **Supplementary Tables 1 and 2**. The slide-mounted tissue section was placed on the stage of an inverted microscope (Nikon, Ti-E) and a custom gasket was then clamped onto the slide, allowing the tissue to be submerged in 1.5 mL of buffer solution (TBST, protein assay; 2X SSC +0.1% Tween 20, RNA assay). The gasket clamp design allowed the buffer to be accessed from above by a microcapillary tip (100 µm inner diameter). The microcapillary tip was connected to a syringe pump (Cavro XCalibur) primed with buffer solution, allowing for accurate aspiration of small volumes (<2.5 µL). Additionally, the tip was mounted to a separate, vertically-aligned z-stage (ASI LS-50), which provided sub-micron tip position accuracy over the tissue. Under the microscope, wide field fluorescence imaging was performed with epi-illumination from visible LED light engine (Lumencor, SOLA). The tissue area of interest was then located using fluorescence imaging with a 4x objective (Nikon). This was followed by 20x (Nikon, ELWD) fluorescence scanning. Each 20x image corresponds to 665µm × 665µm of tissue area with a CMOS camera (Hamamatsu, Flash 4.0). The 20x images were assembled to yield a high-resolution image of the tissue area of interest. The specific regions of interest (ROIs) for molecular profiling were then selected based on the fluorescence information and sequentially processed by the microscope automation. For the data presented in **Fig. 2g, h**, the same overall workflow (as above) is accomplished with a custom-built high-speed automated system (GeoMx Digital Spatial Profiler, NanoString, Seattle WA) and an integrated instrument software. Also, the single-DMD configuration (custom design by NanoString) was enhanced to utilize a dual-DMD configuration, with two semiconductor chips operated in series, in order to provide enhanced background signal reduction.

The steps performed for each ROI by the microscope automation were as follows: First, the microcapillary tip was washed by dispensing clean buffer out the capillary and into a wash station. Next, the tissue slide was washed by exchanging the buffer solution on the slide via the inlet and outlet wash ports on the gasket clamp. The microcapillary tip was then moved into position 50 µm above the ROI. The local area of tissue around the ROI was washed by dispensing 100 µL of buffer solution from the microcapillary. Then, the ROI was selectively illuminated with UV light to release the indexing oligos by coupling UV LED light with a digital mirror device (DMD) module (Andor, Mosaics3). The DDMD configuration used in **Fig. 2g, h** is a custom optical module built directly from a Texas Instrument DMD chip. UV LED light was collimated to be reflected from the DMD surface into the microscope objective, and focused at the sample tissue (365nm, ~125mW/Cm2 or 385nm, ~800mW/Cm2; 1-10 second exposure). Each micro mirror unit in the DMD corresponds to ~1µm^2^ area of sample and reflects the UV light in a controlled pattern based on the ROl selection in the image. Following each UV illumination cycle, the eluent was collected from the local region via microcapillary aspiration and transferred to an individual well of a microtiter plate. Once all ROIs were processed, pools of released indexing oligos were hybridized to NanoString optical barcodes for digital counting and subsequently analyzed with an nCounter Analysis System or NGS readout using the protocols below.

### Custom UV illumination mask creation

Custom masks were created to define custom regions of interest for UV illumination for experiments described in **Fig. 7** and **Fig. 8**. These custom masks were used by the DMD device (described above) to determine which mirrors would be utilized to direct UV light to these custom regions. For each region of interest, 20x stacked tiff images of four fluorescent images were processed with ImageJ^26^ to create these custom masks. For tumor and tumor inverse segments (**Fig. 7a-d**), each stacked tiff was split into four separate images for custom thresholding. The tumor marker, PanCK, image was thresholded manually to match the PanCK staining pattern and converted to a binary mask. To fill holes, a “fill holes” secondary mask was generated using Analyze Particles with settings *size* = 0-700 pixels, *circularity* = 0.35-1.00, and *show* = masks. A selection was generated on the “fill holes” secondary mask, copied onto the tumor mask and the holes were filled to generate a tumor_filled-holes mask. To remove small particles, a “remove particles” secondary mask was generated by inverting the tumor_filled-holes mask and using “Analyze Particles” with settings *size* = 0-60 *circularity* = 0-1.00, and *show* = masks. A selection was generated on the “remove particles” secondary mask, copied onto the tumor_filled-holes mask, filled holes, removed selection, and inverted the mask to generate a tumor_filled-hole_removed-particles mask that will be referred to as the Original_Tumor_Mask. The Original_Tumor_Mask was inverted, eroded 3 times, removed small particles, created a centered 665 µm diameter circle selection, and cleared outside to generate the final tumor segment. The tumor inverse mask was generated by inverting the Original_Tumor_Mask, dilated 4 times, created a centered 665 µm diameter circle selection, and cleared outside to generate the final tumor inverse segment.

For rare cell profiling of macrophages (**Fig. 7e-f**), each stacked tiff was split into four separate images. The macrophage marker, CD68, image was thresholded manually to match the CD68 staining pattern, converted to a binary mask and dilated 2 times. To fill holes, a “fill holes” secondary mask was generated using Analyze Particles with settings *size* = 0-1000 pixels, *circularity* = 0.30-1.00, and *show* = masks. A selection was generated on the “fill holes” secondary mask, copied onto the CD68 mask and the holes were filled to generate a CD68_filled-holes mask. To remove small particles, a “remove particles” secondary mask was generated by inverting the CD68_filled-holes mask, eroding 2 times, and using “Analyze Particles” with settings *size* = 0-60, *circularity* = 0-1.00, and *show* = masks. A selection was generated on the “remove particles” secondary mask, copied onto the CD68_filled-holes mask, holes filled, removed selection, and mask inverted to generate a the final CD68 segment.

For the contour profiling of macrophage enriched regions (**Fig. 8)**, each stacked tiff was split into four images and the macrophage marker, CD68, image was used for contour profiling. The polygon selection tool was used to trace along the center of the macrophage region and the selection was enlarged on each side by 25 µm to create the 50 µm width macrophage enriched segment. The macrophage enriched segment was then serially enlarged by 50 µm to create radiating 50 µm thick segments away from the macrophage-enriched segment. The wand tool was then used to move each 50 µm segment to their respective 665 × 665 µm mask.

### Sample preparation for protein profiling and IHC

Each selected primary antibody was coupled to a unique 70 nt indexing oligo (NanoString, custom conjugation service). All assays were performed on 5 µm FFPE sections mounted onto charged slides. Deparaffinization and rehydration of tissue was performed by incubating slides in 3 washes of CitriSolv (Decon Labs, Inc., 1601) for 5 minutes each, 2 washes of 100% ethanol for 10 minutes each, 2 washes of 95% ethanol for 10 minutes each, and 2 washes of dH_2_O for 5 minutes each. For antigen retrieval, slides were then placed in a plastic Coplin jar containing 1X Citrate Buffer pH 6.0 (Sigma, C9999) and covered with a lid. The Coplin jar was placed into a pressure cooker (BioSB, BSB7008) and run on high pressure and temperature for 15 minutes. The Coplin jar was removed from the pressure cooker and cooled at room temperature for 25 minutes. Slides were washed with 5 changes of 1X TBS-T (Cell Signaling Technology, 9997) for 2 minutes each. Excess TBS-T was removed from the slide, and a hydrophobic barrier was drawn around each tissue section with a hydrophobic pen (Vector Laboratories, H-4000). Slides were then incubated with blocking buffer [1X TBS-T, 5% Goat Serum (Sigma-Aldrich, G9023-5ML), 0.1 mg/mL salmon sperm DNA (Sigma-Aldrich, D7656), and 10 mg/mL dextran sulfate (Sigma-Aldrich, 67578-5G)] for 1 hour. Slides were washed with 3 changes of 1X TBS-T for 2 minutes each. Primary antibodies were diluted in antibody diluent [Signal Stain Antibody Diluent (Cell Signaling Technology, 8112), 0.1 mg/mL salmon sperm DNA, and 10 mg/mL dextran sulfate]. Tissue sections were then covered with diluted primary antibody solution. Slides were incubated at 4°C in a humidity chamber overnight. Primary antibody was aspirated from slides and washed with 3 changes of 1X TBS-T for 10 minutes each. DNA was counterstained with 100 nM SYTO 83 (ThermoFisher, S11364) or 500 nM SYTO 13 (ThermoFisher, S7575) in 1X TBS-T for 15 minutes. Excess DNA counterstain was removed with 5 changes TBS-T, and slide was processed in an automated fashion on the instrument described above.

### RNA profiling probe design

DNA oligo probes were designed to bind mRNA targets. From 5’ to 3’, they each comprised a 35-40 nt target complementary sequence, an iSpPC UV photocleavable linker (Integrated DNA Technologies), and a 60 nt indexing oligo sequence. All RNA detection probes and indexing oligo sequence complements were ion-exchange HPLC purified (Integrated DNA Technologies). Up to 10 oligo RNA detection probes were designed per target mRNA.

### Sample preparation for RNA profiling

To perform *in situ* hybridization, 5 µm FFPE tissue sections mounted on positively charged histology slides were processed on a Leica Bond Rxm system. Sections were baked, deparaffinized, rehydrated in ethanol, and washed in PBS. Targets were retrieved for 10 to 20 minutes in 1X Tris-EDTA pH 9.0 buffer (Sigma Aldrich, SRE0063) at 85°C (tonsil and cell pellet arrays) or 100°C (colorectal samples). Next, tissues were washed in PBS, incubated in 1 µg/mL proteinase K (Thermo Fisher Scientific, AM2546) in PBS for 5 to 25 minutes at 37°C and washed again in PBS. Tissues were removed from the Leica Bond and incubated overnight at 37°C with hybridization solution (20 nM RNA detection probes per target; 100 μg/mL denatured, sonicated salmon sperm DNA (Sigma-Aldrich, D7656); 2.5% dextran sulfate (Sigma-Aldrich, 67578-5G); 0.2% BSA (Sigma, A1933); 40% deionized formamide (Thermo Fisher Scientific, AM9344); 2X SSC (Sigma, S6639)). During incubation, slides were covered with HybriSlip Hybridization Covers (Grace BioLabs, 714022). Following incubation, HybriSlip covers were gently removed by soaking in 2X SSC + 0.1% Tween 20. Two 25-minute stringent washes were performed in 50% formamide in 2X SSC at 37°C. Tissues were washed for 5 minutes in 2X SSC then incubated in blocking buffer (composition above) for 30 minutes at room temperature in a humidity chamber. 100uM SYTO 13 and fluorescently-conjugated antibodies targeting PanCK and CD3 in blocking buffer were applied to each section for 1 hour at room temperature, then washed for 5 minutes in fresh 2X SSC. Indexing oligos were then cleaved and quantitated as previously described.

### nCounter hybridization assay for photocleaved oligo counting

Hybridization of cleaved indexing oligos to fluorescent barcodes was performed using the nCounter Protein TagSet reagents. Indexing oligos were denatured at 95°C for 3 to 5 minutes and placed on ice for 2 minutes. A master mix was created by adding 70 μL of hybridization buffer to the Protein TagSet tube. A 7 μL aliquot of master mix was added to each of 12 hybridization tubes. Depending on the experiment, 2 to 8 µL denatured protein sample was added to each tube and each hybridization was brought to a final volume of 15 μL with DEPC treated water. Hybridizations were performed at 65°C overnight in a thermocycler. After hybridization, samples were processed using the nCounter Prep Station and Digital Analyzer as per manufacturer instructions.

### RNAscope *in situ* hybridization

RNAscope in situ hybridization was performed on 5 µm FFPE tissue sections using RNAscope Multiplex Fluorescent Reagent Kit (ACD, 322800) as per manufacturer’s instructions. Epitope retrieval was performed for 15 minutes at 88°C and proteinase K digestions was performed at 40°C for 15 minutes. Probes used include CD3 (ACD 553978), CD20 (ACD 426778), and CD40 (ACD 578478) and were visualized using either TSA Plus Cyanine 3 (Perkin Elmer, NEL744001KT) or TSA Plus Cyanine 5 System (NEL745001KT).

All cell pellet slides were fluorescently imaged using the Nikon Eclipse TE2000-E with a 40X objective. Images were captured with Nikon Elements commercial software. For imaging, two Z-stacks, from the top to bottom focal planes, of each cell pellet. Maximum Z projection images were created with Nikon Elements software across all channels. QuPath software (https://qupath.github.io/) was used to quantify the number of RNAscope spots imaged per FOV. For this, scripts were run for each FOV which calculated the numbers of cells, through nuclei counting in the DAPI channel, and number of fluorescent spots from the RNAscope assay. Nuclei counting did not appear accurate for some cell types which had nuclei that could not be readily distinguished, therefore total spots per FOV was reported. Watershed detection of nuclei (‘qupath.imagej.detect.nuclei.WatershedCellDetection’ plugin) was performed with the following settings: “requested Pixel Size Microns”: 0.5, “background Radius Microns”: 8.0, “median Radius Microns”: 0.0, “sigma Microns”: 1.5, “min Area Microns”: 10.0, “max Area Microns”: 400.0, “threshold”: 100.0, “watershed Post Process”: true, “cell Expansion Microns”: 20.0, “include Nuclei”: true, “smooth Boundaries”: true, “make Measurements”: true. RNAscope spot detection (‘qupath.imagej.detect.cells.SubcellularDetection’. plugin) was performed with the following settings: “detection[Channel]ȝ: 300.0, “do Smoothing”: true, “split By Intensity”: true, “split By Shape”: true, “spot Size Microns”: 4.0, “min Spot Size Microns”: 0.2, “max Spot Size Microns”: 25.0, “include Clustersȝ: true, where the detection channel is the corresponding channel used for RNAscope visualization.

### Cell Pellet Purified RNA Quantification on nCounter

RNA expression for FFPE cell pellets were analyzed to select positive and negative controls for IHC (RNA expression data not shown). For this, deparaffinization and rehydration of FFPE cell pellets was performed by incubating slides in 2 washes of CitriSolv (Decon Labs, Inc., 1601) for 2 minutes each and in 100% ethanol for 2 minutes. The slides were air dried then dipped in 3% glycerol (MP Bio, 3055-034). Excess 3% glycerol was removed from slide and the cell pellet was scraped in a single direction on the slide with a clean razor then transferred to a microcentrifuge tube. Lysis buffer mix consisting of 100 μL Lysis buffer (Vial 1, Roche FFPET Isolation Kit # 06650775001), 100 μL dH2O, and 200 μL of 20 mg/mL Proteinase K (Roche FFPET Isolation Kit # 06650775001) was added to each tube. Tubes were mixed and samples spun down then incubated at 55°C for 30 minutes while shaking at 600 RPM. RNA hybridization was performed using the nCounter PanCancer IO 360 Gene Expression Panel. Target lists for this panel nCounter panels can be found on NanoString’s website. For the assay, a master mix was created by adding 70 μL of hybridization buffer to the Reporter Probe tube. 8 μL aliquots of the master mix was added to each of the 12 hybridization tubes. 5 μL of RNA samples (100 ng total RNA) were added to each tube followed by 2 μl of the Capture Probe set. Tubes were mixed and samples spun down. Hybridizations were performed at 65°C overnight in a thermocycler with a 70°C heated lid. After hybridization, samples were processed using the nCounter PrepStation and Digital Analyzer as per manufacture instructions. Digital counts from barcodes corresponding to protein probes were normalized to ERCC counts and housekeeping gene counts.

### Tissue Immunohistochemistry with DAB-based secondary detection

Immunohistochemistry of FFPE tissues (**Fig. 2b, Supplementary Fig. 5f**) was performed using the Leica Biosystems Bond Rx. Pretreatment of FFPE slides consisted of deparaffinization for 30 minutes at 72°C using Bond Dewax solution followed by antigen retrieval for 20 minutes using the Bond ER #1 solution. Slides were blocked for 1 hour with blocking buffer [1X TBS-T, 5% Goat Serum (Sigma-Aldrich, G9023-5ML), 0.1 mg/mL salmon sperm DNA (Sigma-Aldrich, D7656), and 10 mg/mL dextran sulfate (Sigma-Aldrich, 67578-5G)] and then incubated with the primary antibody for 60 minutes. The primary antibody was diluted (see antibody table for concentrations) in antibody diluent [Signal Stain Antibody Diluent (Cell Signaling Technology, 8112), 0.1 mg/mL salmon sperm DNA, and 10 mg/mL dextran sulfate]. Secondary detection was performed this the Bond Polymer Refine Detection Kit (Leica, DS9800). Tissue sections were imaged using a Nikon Eclipse Ci light microscope with 4x or 20x objectives and Nikon Elements software.

### NGS readout for photocleaved oligo counting

Each ROI was uniquely indexed using Illumina’s i5 × i7 dual-indexing system. 4 uL of the aspirate containing the photocleaved oligos was used in a PCR reaction with 1 uM of i5 primer, 1 uM i7 primer, and 1X NSTG PCR Master Mix. Thermocycler conditions were 37°C for 30 min, 50°C for 10 min, 95°C for 3 min, 18 cycles of 95°C for 15 sec, 65°C for 60 sec, 68°C for 30 sec, and final extension of 68°C for 5 min. PCR reactions were purified with two rounds of AMPure XP beads (Beckman Coulter) at 1.2x bead-to-sample ratio. Libraries were paired-end sequenced (2×75) on a MiSeq with 25.9M total aligned reads and a median of 0.2M aligned reads per ROI (range 1,128 to 1.1 M). Each library (or ROI) was demultiplex on-instrument to produce one FASTQ file per ROI. Raw sequencing reads were processed for high quality using Trim Galore and Paired-End reAd mergeR. Reads were then aligned to analyte barcode with Bowtie. For NGS digital counts, PCR duplicates were removed using UMI-tools with a hamming distance set at three.

### Data Analysis and Visualization

For nCounter data analysis, digital counts from barcodes corresponding to protein or RNA probes were first normalized with internal spike-in controls (ERCCs) to account for system variation. Box plots **(Fig. 2d, Supplementary Fig. 1)** were created with the ggplot2 R package. The upper whisker extends from the hinge to the highest value that is within 1.5 * IQR of the hinge, where IQR is the inter-quartile range, or distance between the first and third quartiles. The lower whisker extends from the hinge to the lowest value within 1.5 * IQR of the hinge. Data beyond the end of the whiskers are outliers and plotted as points. IO360 nCounter counts from 100 ng purified RNA (see methods) were used to classify which cell lines are likely positive (> 100 counts) or negative (<10 counts) for protein expression. The conservative threshold of 10 counts for a negative cell line was selected since this approach will likely be less accurate to estimate true positives, as compared to true negatives. The true negative will also likely be more accurately called because RNA is necessary, but not sufficient, for protein expression. Since there is high confidence in calling these true negatives with this approach, limits-of-detection (LODs) were estimated with the negative cell lines in each target/ROI condition. For this, the average plus 3 times the standard deviation was used to estimate the LOD of each antibody, for this experiment. Targets with nCounter counts between 10 and 100, targets expected to be expressed in all cell lines (housekeeper), or antibodies with no specificity (IgG controls) were labeled as “Intermediate” or “Unknown” expression. Dot plots **(Fig. 2f, 2h, 3d, 3e Supplementary Fig. 2b, 2d, 4a, 4b, 5b)** were created and R2 analysis performed with Excel (Microsoft) or were created with the ggplot2 R package **(Fig. 4c, 4d, 4e).** Clustered heatmaps **(Fig. 2e, 5d, 5d, 5g, 5h, 6d, 7c, 8c, Supplementary Fig. 5d.)** were created with the pheatmap R package. Data was log10 transformed (log10 baseR function) and scaled (scale baseR function) for each target prior to clustering. Clustering was with the pheatmap “correlation” function. For **Fig. 2e**, each ROI was first normalized with the average of the geomean of all targets and for **Fig. 7c, 8c** each ROI was first normalized by area to normalize for differences in ROI sizes. Gridded heatmaps **(Fig. 6b, 6c, 6h, 6i, Supplementary Fig. 6)** were created with the heatmap function in the Seaborn Python package. TMA heatmaps **(Fig. 2c, Supplementary Fig. 2c, 5e)** were created with the ggplot2 R package.

## Data availability

The data that support the findings of this study are available from the authors on reasonable request, see author contributions for specific data sets.

## Availability of Materials

All unique materials used are readily available from Nanostring Technologies, Inc.

## Supporting information

Fig. 1 - hi res

Fig. 2 - hi res

Fig. 3 - hi res

Fig. 4 - hi res

Fig. 5 - hi res

Fig. 6 - hi res

Fig. 7 - hi res

Fig. 8 - hi res

Supplemental Fig. 1 - hi res

Supplemental Fig. 2 - hi res

Supplemental Fig. 3 - hi res

Supplemental Fig. 4 - hi res

Supplemental Fig. 5 - hi res

Supplemental Fig. 6 - hi res

## Acknowledgements

Philippa Webster for preliminary protocol development. Alison VanSchoiack for editing the manuscript. Heather Metz for performing IHC.

## Author contributions

C.R.M., S.E.C, and J.M.B wrote the manuscript, C.M. prepared the figures, J.M.B. and G.M. conceived the project., J.M.B., D.L.D., and C.R.M. supervised the project., C.R.M. supervised experimental studies., G.G. supervised antibody labeling, C.R.M., G.T.O., S.E.C., D.Z., K.S., K.B., and J.M.B designed experiments. K.B. performed antibody validation experiments. G.T.O. and K.B. performed protein profiling experiments. Y.L. and S.E.C. performed IHC interpretation, C.M. developed the immunohistochemistry protocol, C.M. and K.S. developed the RNA profiling protocol, D.Z., J.M.-F., performed RNA profiling experiments, K.N. and K.S. performed RNAscope experiments, C.R.M, G.T.O., K.S., and K.B. performed data analysis, J.J, I.B.S. and D.L.D developed initial instrumentation setup, J.J configured the microscope and performed preliminary spatial protein profiling experiments, C.W. and I.B.S. developed the final instrumentation and fluidics setup, I.B.S. developed the instrumentation automation.

---

**Suppl. Fig. S1:**
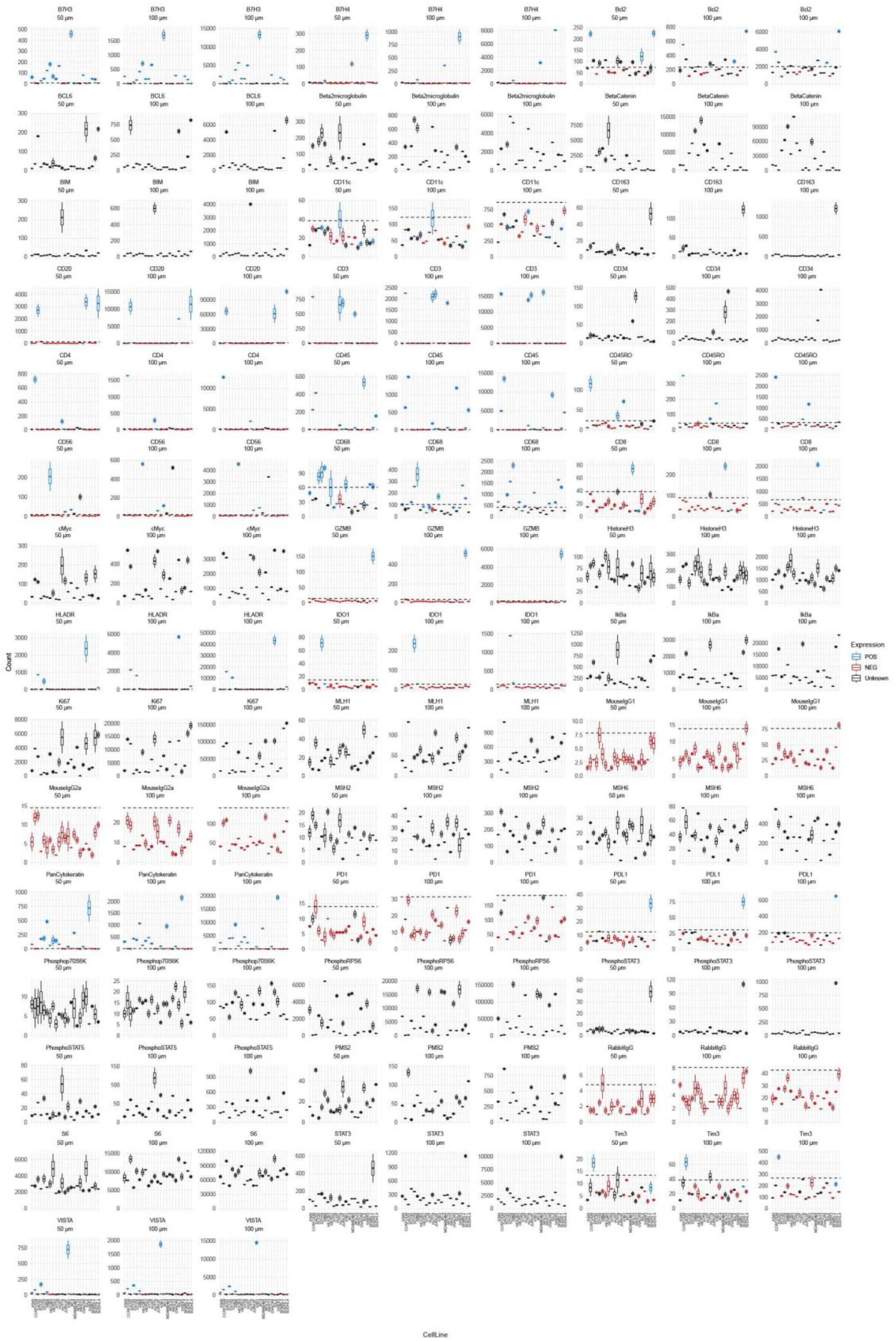
Validation of DSP with oligo-conjugated antibodies. Additional data from experiment presented in **Fig. 2 c,d,e**. All replicate data points (n =2) for all ROI sizes (50 µm, 100 µm, 300 µm) displayed. Antibodies listed in alphabetical order. Y-axes are scaled according the expression range for each antibody and each ROI size. nCounter counts from 100 ng purified RNA (see methods) were used to estimate which cell lines are positive (> 100 counts) or negative (<10 counts). Targets with nCounter counts between 10 and 100, targets expected to be expressed in all cell lines (housekeeper), or antibodies with no specificity (IgG controls) were labeled as “Unknown” expression.

**Suppl. Fig. S2:**
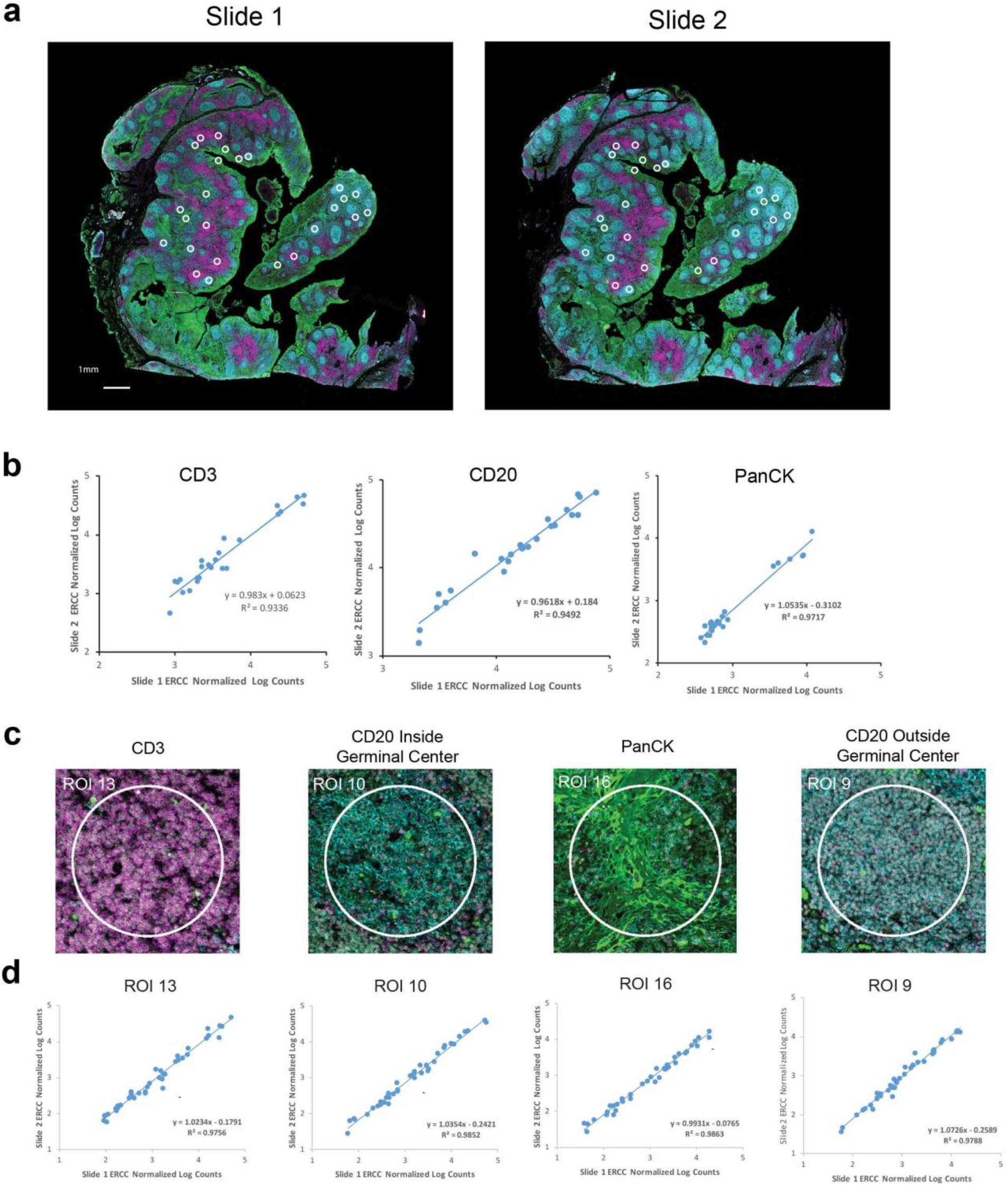
Reproducibility of the DSP system. **a,** Serial sections of tonsil tissues analyzed in **Fig. 1f**. 36 ROIs were selected across each of the tissues, and the 25 overlapping ROIs were used for this analysis. Sections were stained with CD3, CD20, and PanCK antibodies directly conjugated to fluorescent dyes. **b,** Correlation of CD3, CD20, and PanCK on both serial sections across the 25 overlapping ROIs. **c,** Four ROI types selected, based on visualization marker expression, for correlation analysis of the 44 antibodies in the cocktail. **d,** Correlation of antibody counts across overlapping ROIs on adjacent sections.

**Suppl. Fig. S3:**
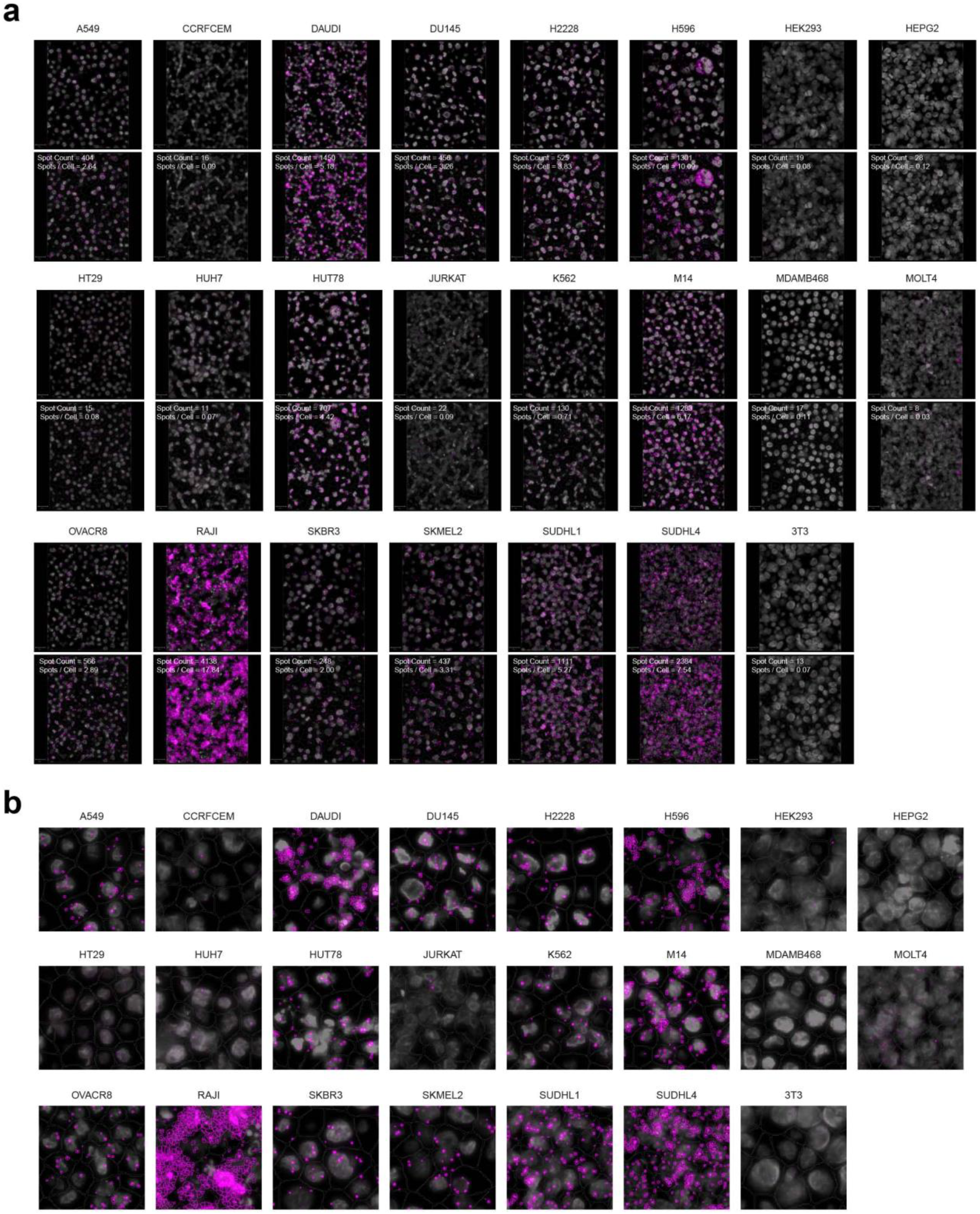
Representative images of CD40 RNAscope quantification. **a,** Representative 40x FOV max intensity Z projection images of CD40 RNAscope quantification. CD40 RNA (magenta) and DAPI (gray). For each cell line, the original image (top) is shown alongside images with estimated cell boundaries from the nuclear stain and RNAscope spot boundaries defined with the QuPath quantification process (bottom). **b,** Closeup of representative quantification boundaries across all cell lines.

**Suppl. Fig. S4:**
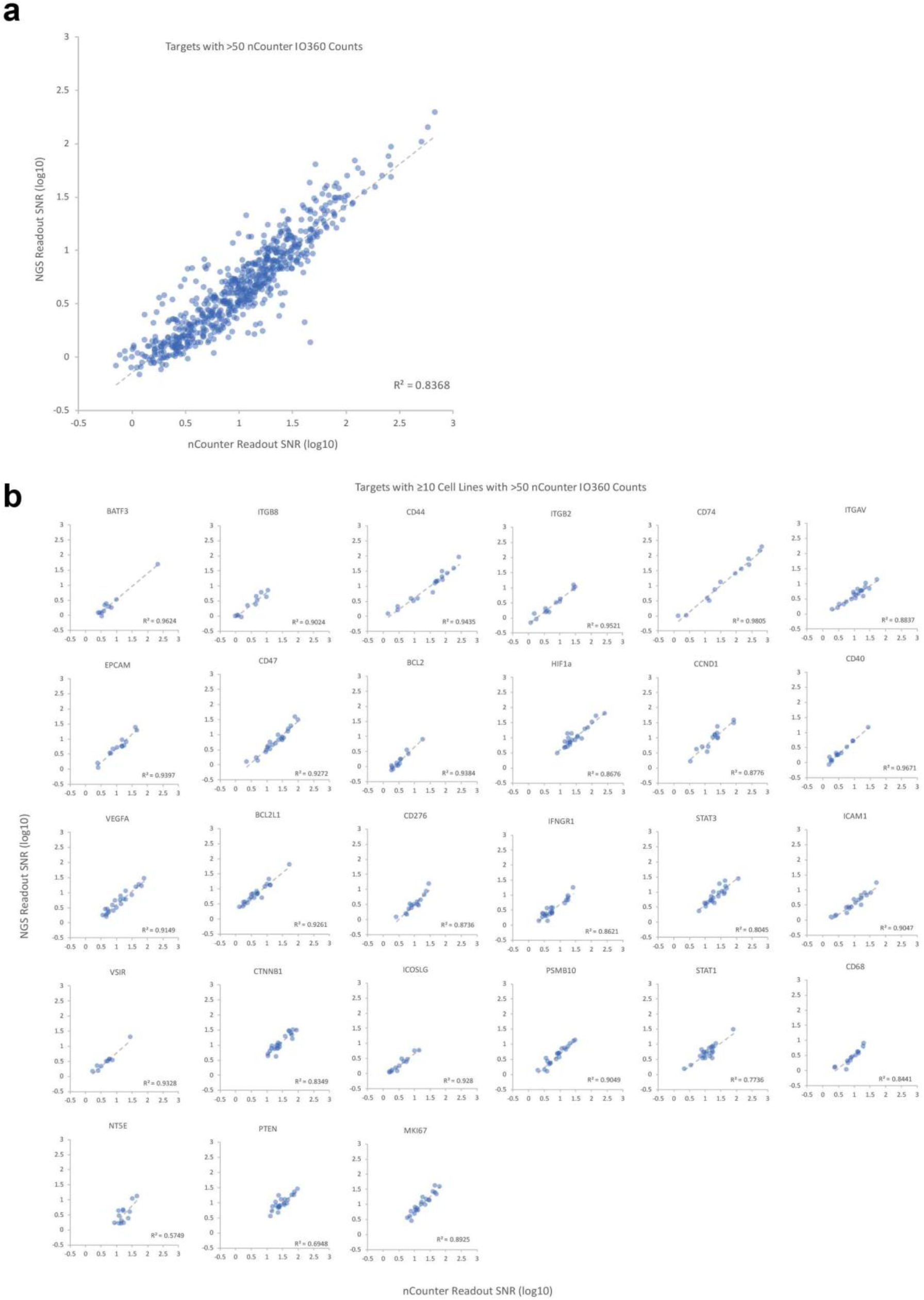
Correlation between nCounter and NGS RNA DSP probes. **a,** nCounter IO360 expression data was used to determine target with expression confidently above background (> 50 counts). These target/cell lines were used to analyze the correlation between the nCounter and NGS RNA DSP probes. Average SNRs were calculated and log transformed. **b,** Individual correlation plots for targets highlight in **Fig. 3e.** Representative targets shown with at least 10 data points significantly above background for nCounter and covering at least one order of magnitude of expression. Individual R^2^ values shown.

**Suppl. Fig. S5:**
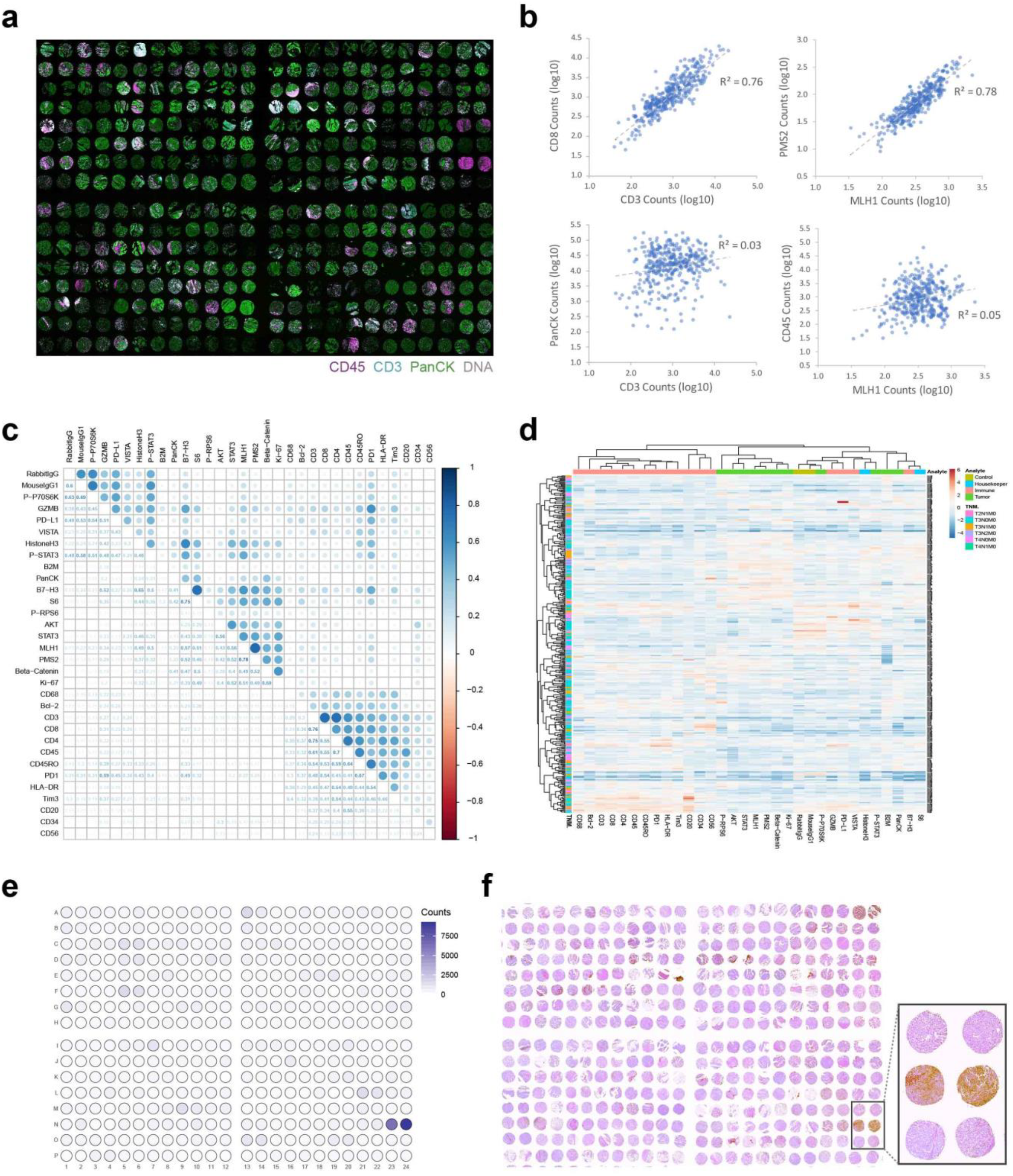
High throughput TMA analysis with geometric ROIs. **a,** Scan of 384 cores analyzed from a high-density colorectal cancer TMA. These 384 cores replicate cores from 192 donor sample. 665 µm diameter circle ROIs where profiled for each core. **b,** Correlation of selected targets across 384 cores. CD3/CD8 (T cell markers) and PMS2/MLH1 (mismatch repair proteins) show high correlation (R^2^ > 0.75), while comparisons of unrelated markers (PanCK/CD3 or CD45/MLH1) show no correlation (R^2^ < 0.05), **c,** Correlation matrix of all target pairs analyzed. **d,** Hierarchically clustered heatmap of region-specific digital data across all protein targets across all 384 cores. **e,** Heatmap of PD-L1 across the 384-cores shows high PD-L1 expression for both replicate cores from a single donor sample. **f,** IHC analysis of PD-L1 on a serial section to confirm the expression of PD-L1 in these two cores.

**Suppl. Fig. S6:**
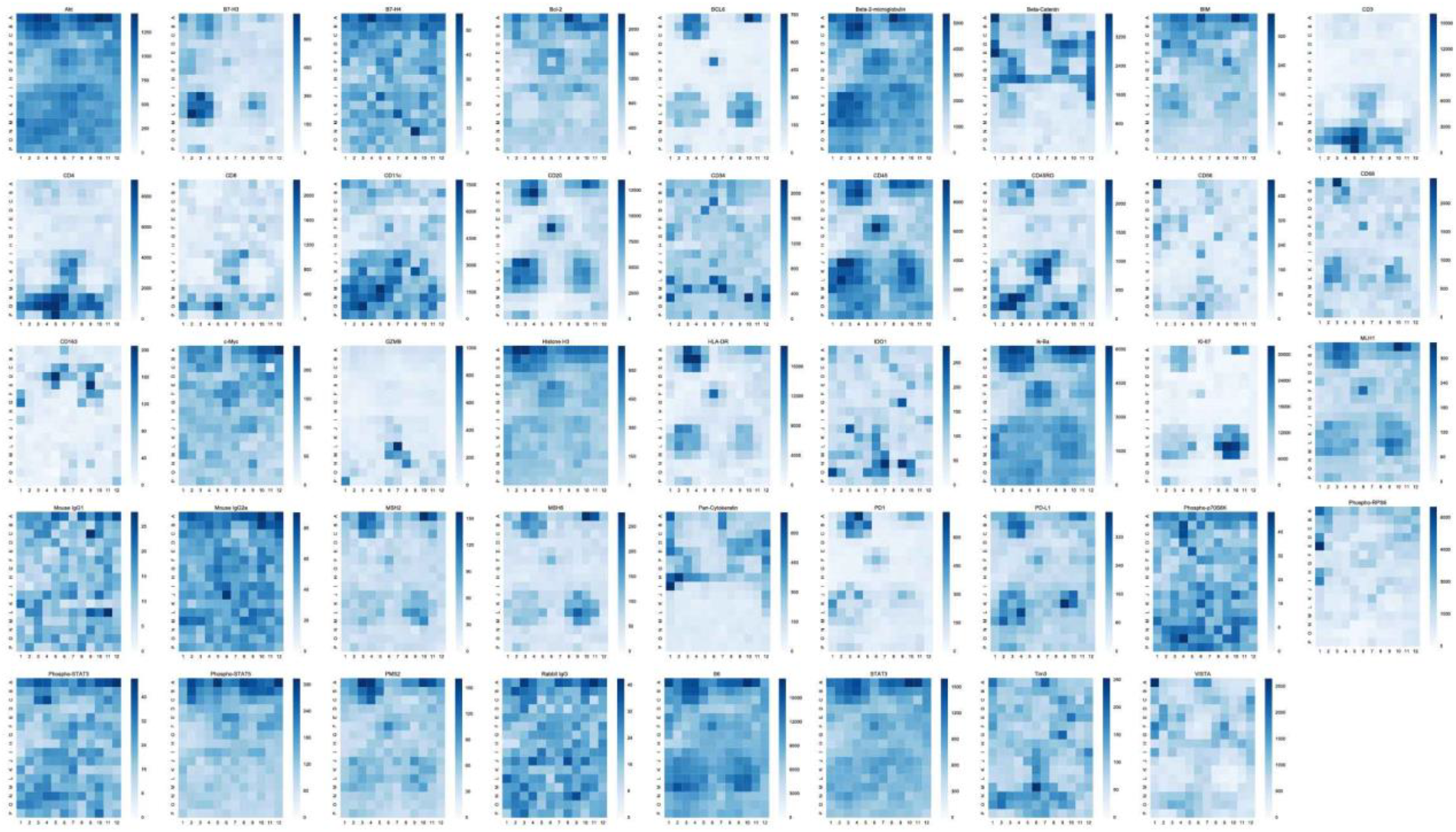
Heatmaps for all protein targets in tonsil. Heatmaps for each antibody analyzed in the gridded analysis presented in **Fig. 6a-f**.

**Suppl. Table S1. Conjugated antibodies used in these studies.** 44 antibodies conjugated to unique photocleavable oligos were mixed into a single cocktail or these studies. NanoString provides a custom antibody number (CAB#) for each unique antibody conjugation clone in lieu of specific identifying information about the antibodies as Nanostring considers this confidential information. The custom antibody number is provided in the supplementary material and will allow investigators who wish to replicate the work to use the same reagents. Conjugated IHC validation (as shown in **Fig. 2b**) was performed for most antibodies in these studies.

**Suppl. Table 2. RNA probes used in these studies.**

Unique RNA probe ID, target transcript, and exon targeted are shown.

