## Supplementary figures and images for "High multiplex, digital spatial profiling of proteins and RNA in fixed tissue using genomic detection methods"

### Fig. 1 - hi res

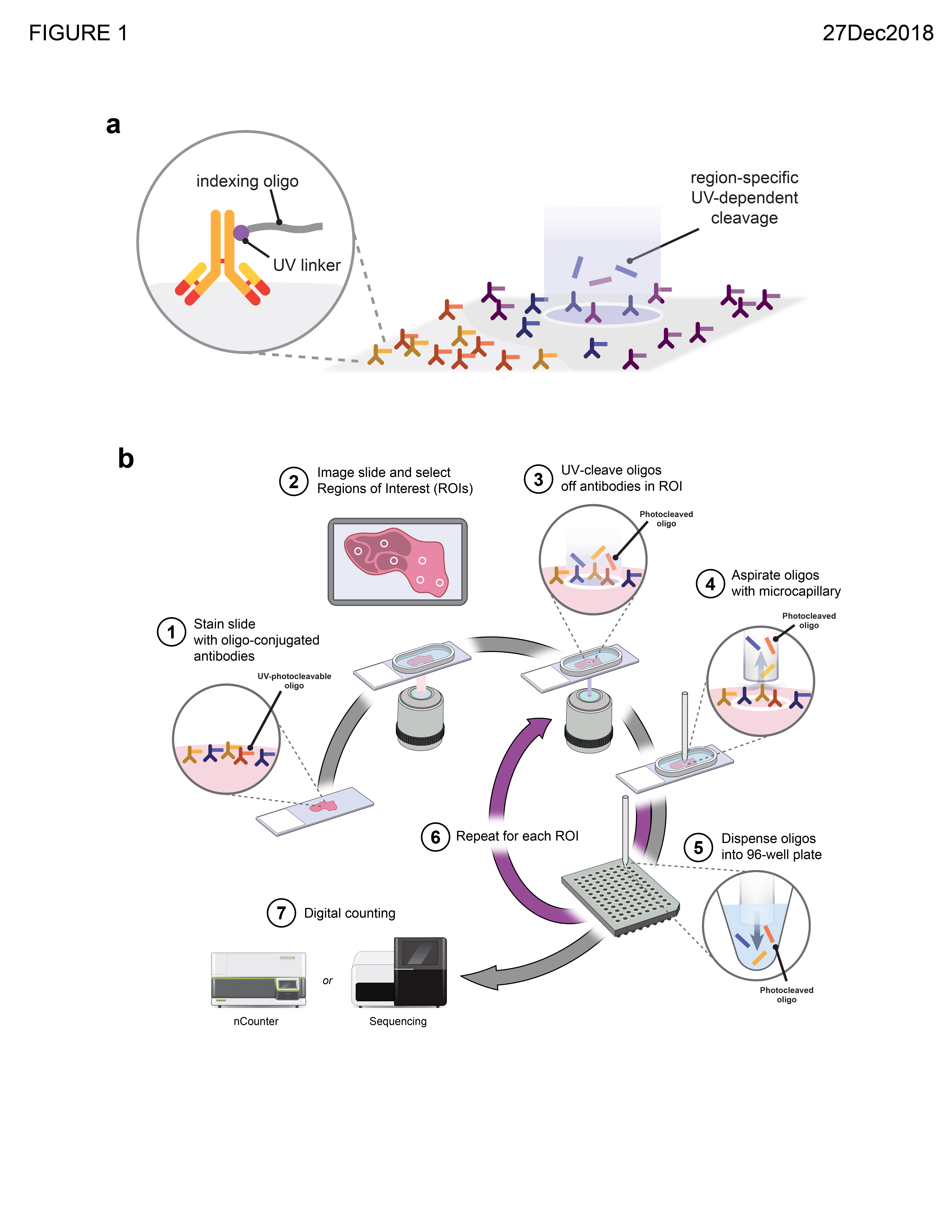

### Fig. 2 - hi res

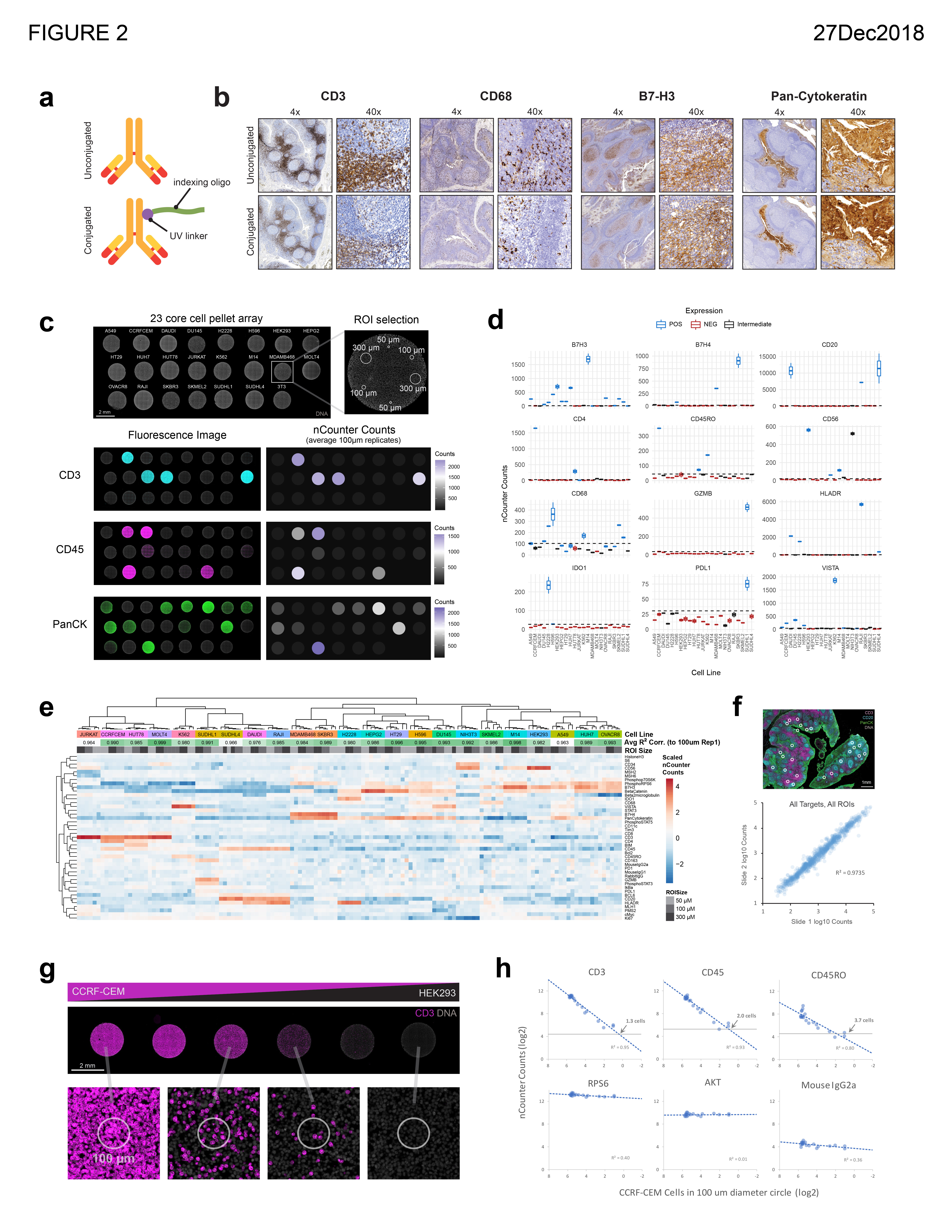

### Fig. 3 - hi res

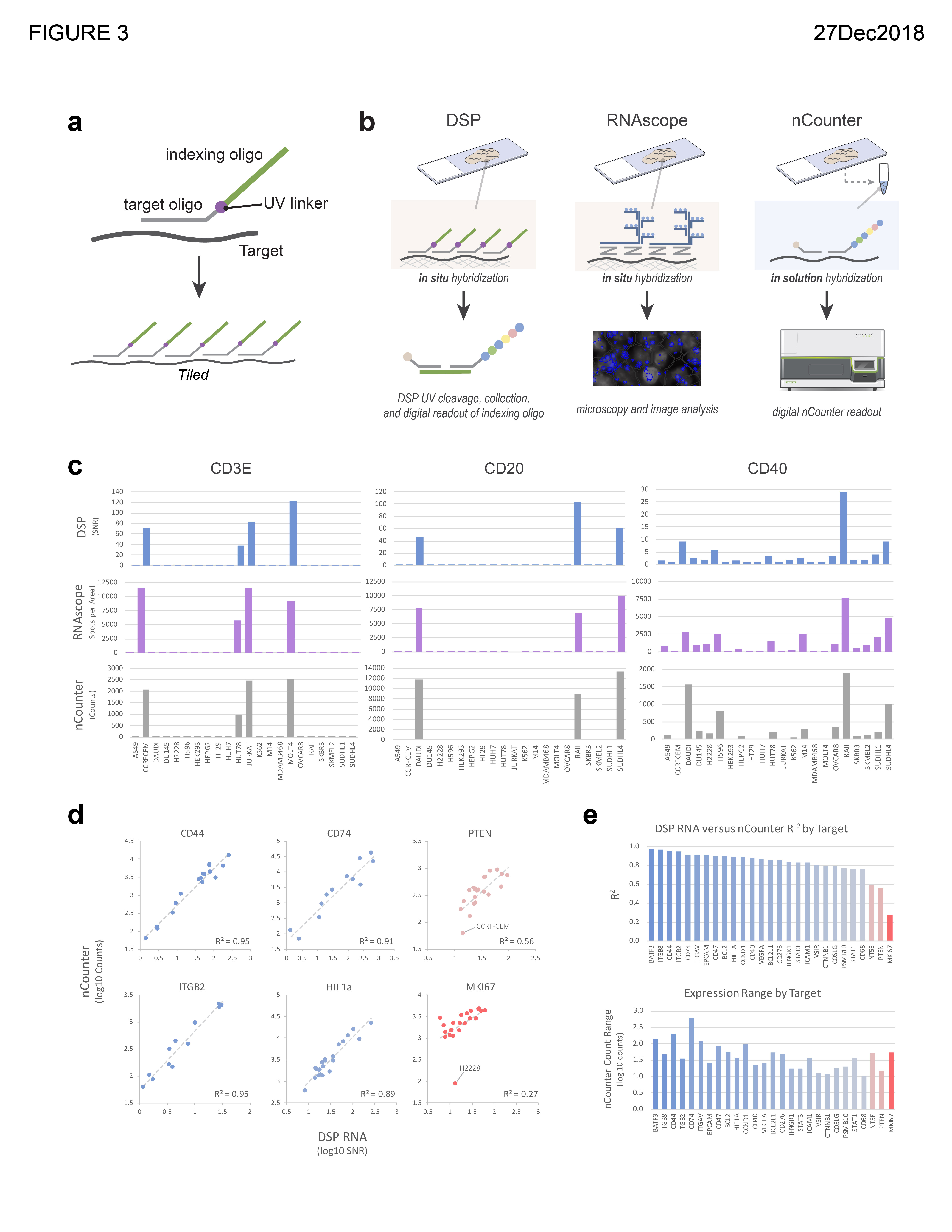

### Fig. 4 - hi res

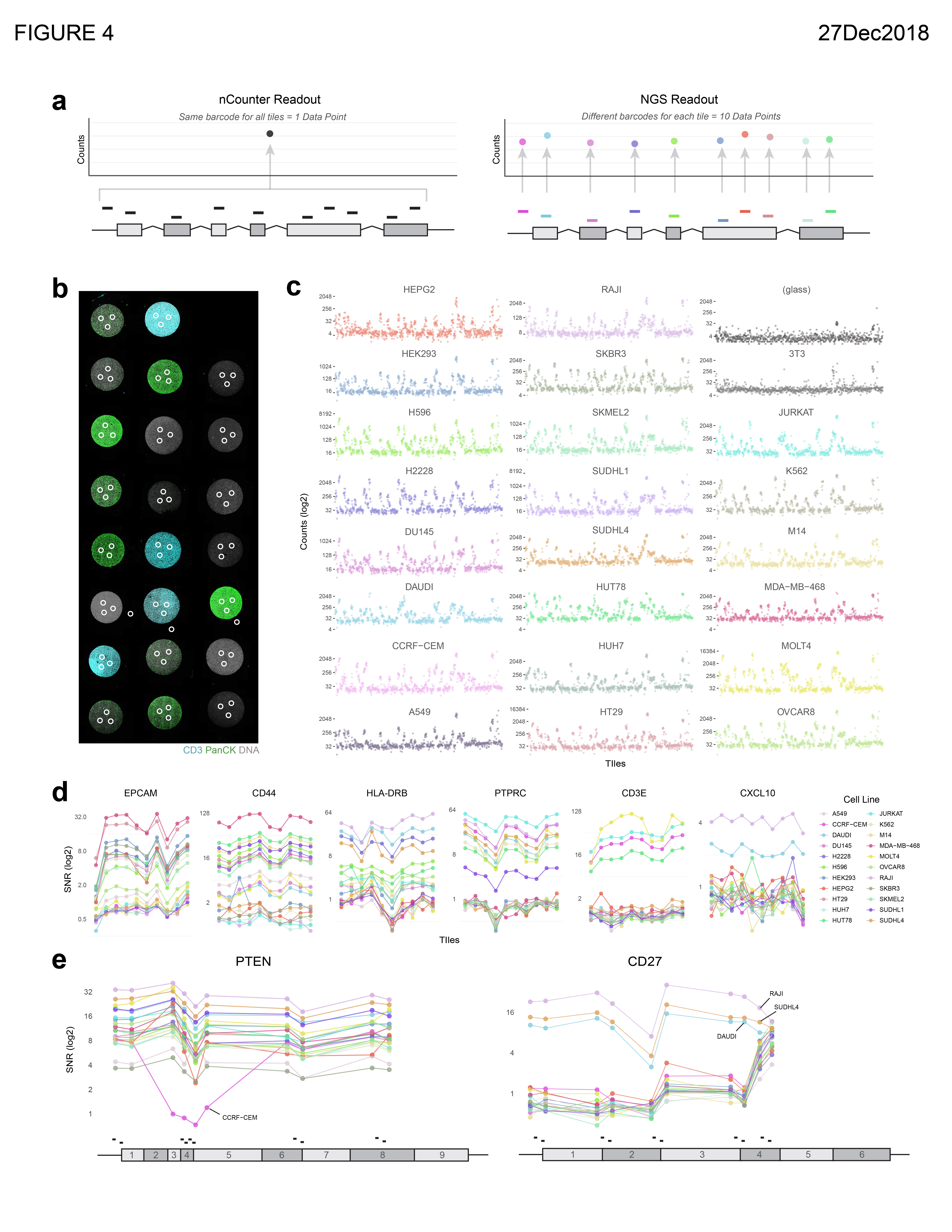

### Fig. 5 - hi res

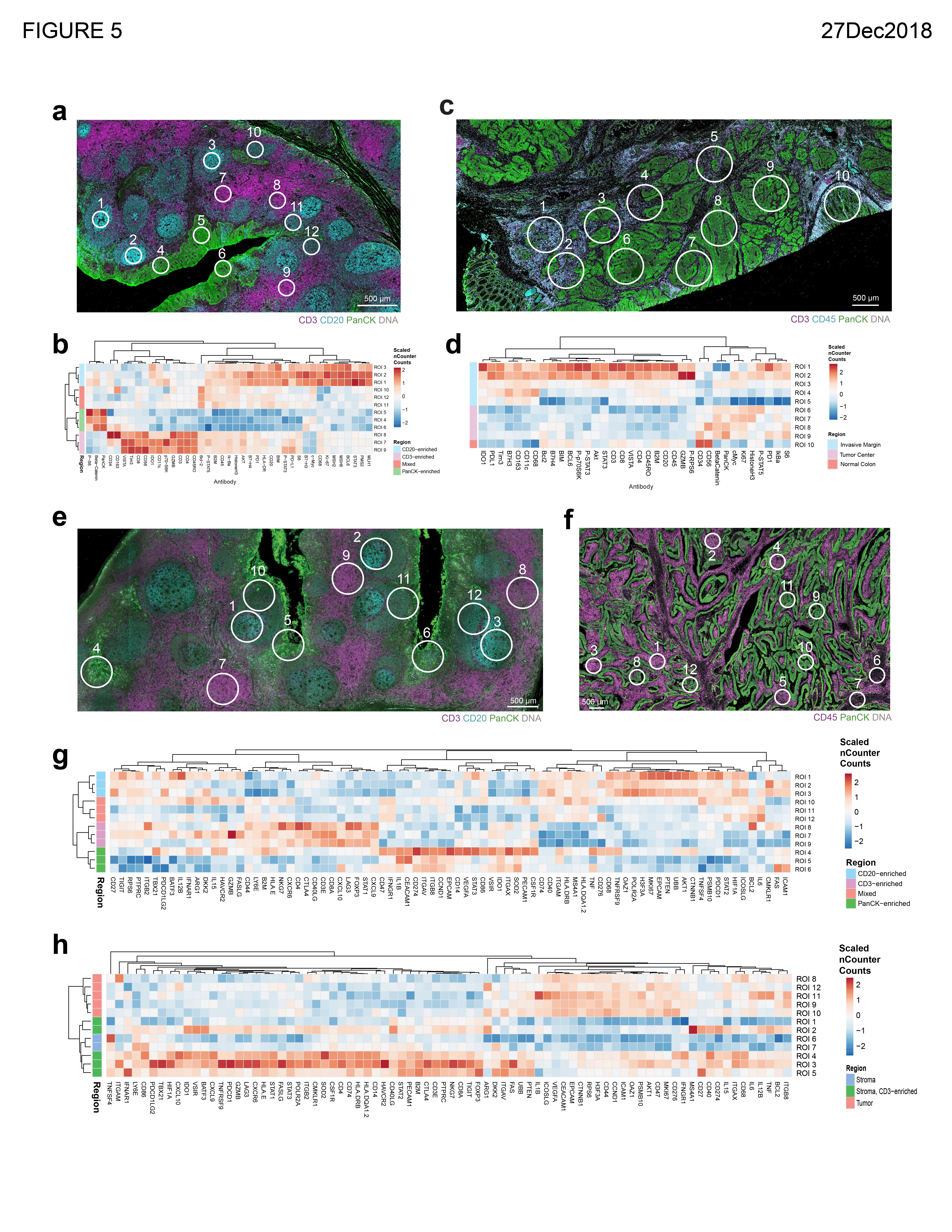

### Fig. 6 - hi res

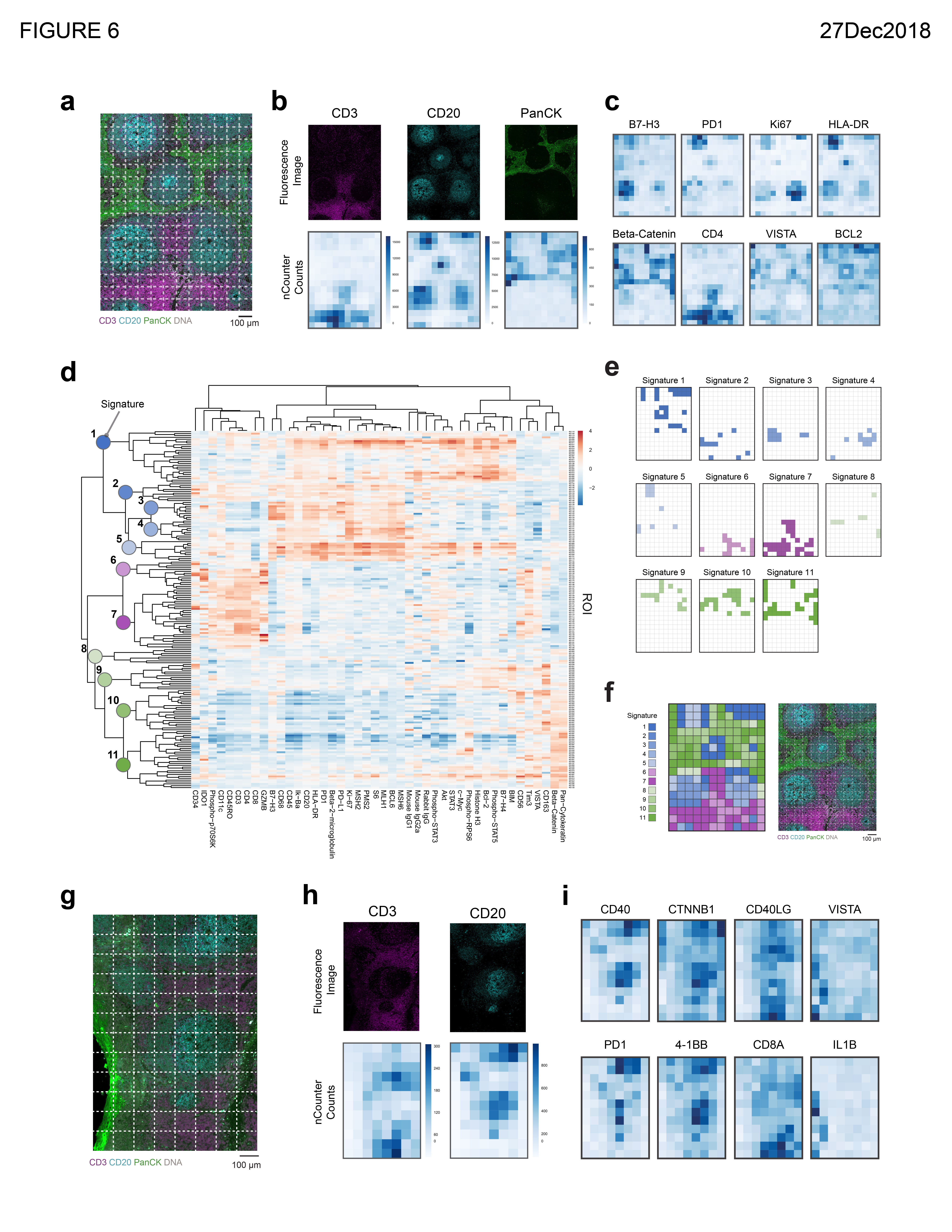

### Fig. 7 - hi res

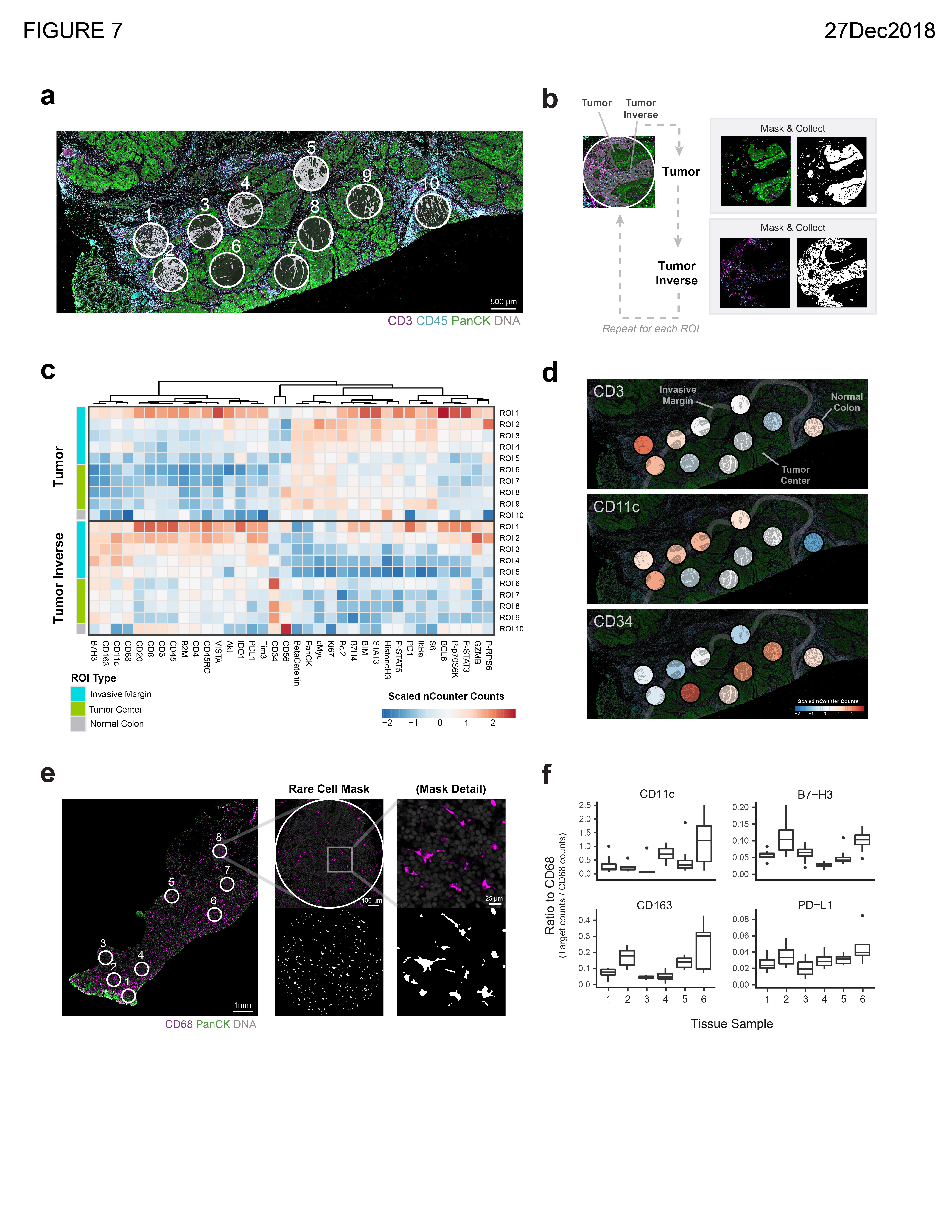

### Fig. 8 - hi res

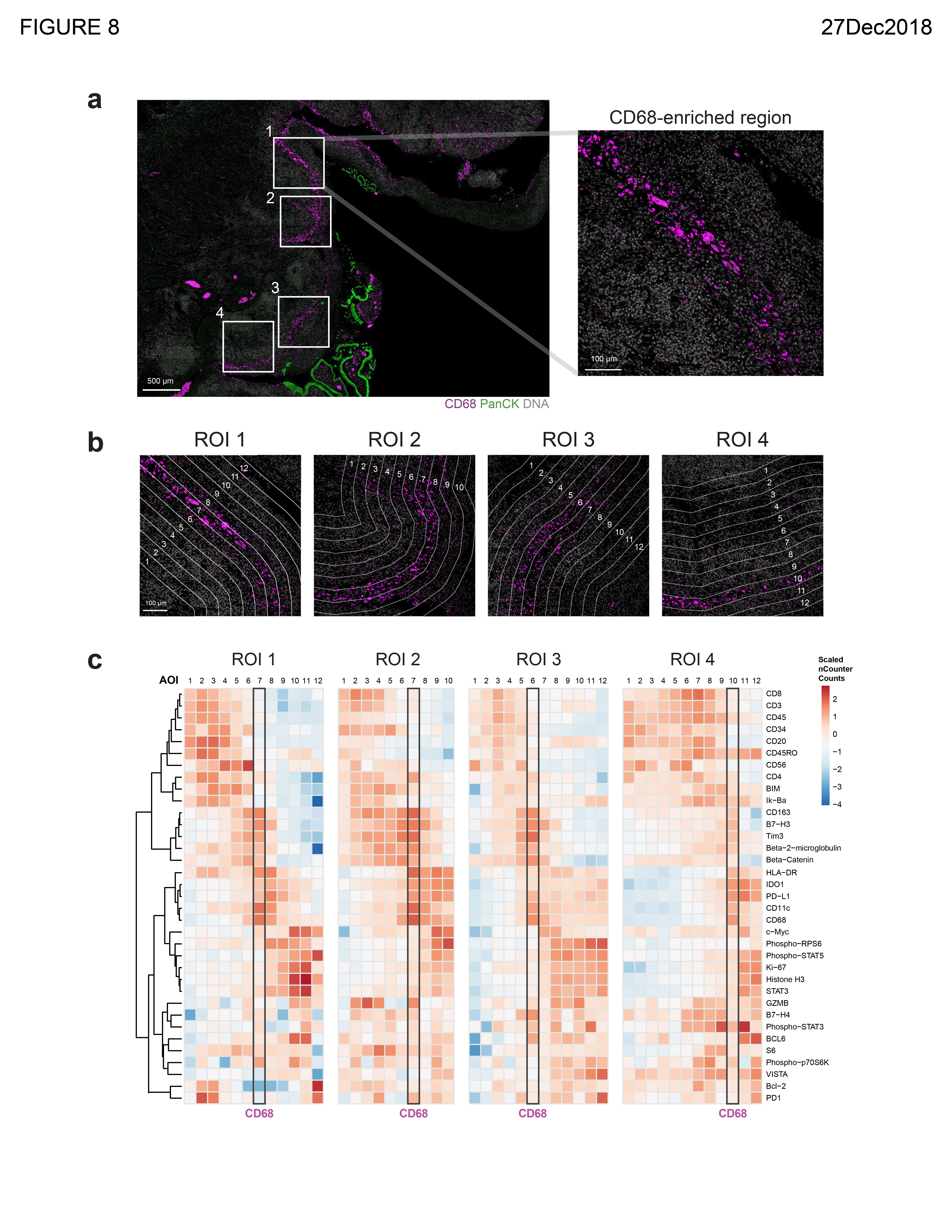

### Supplemental Fig. 1 - hi res

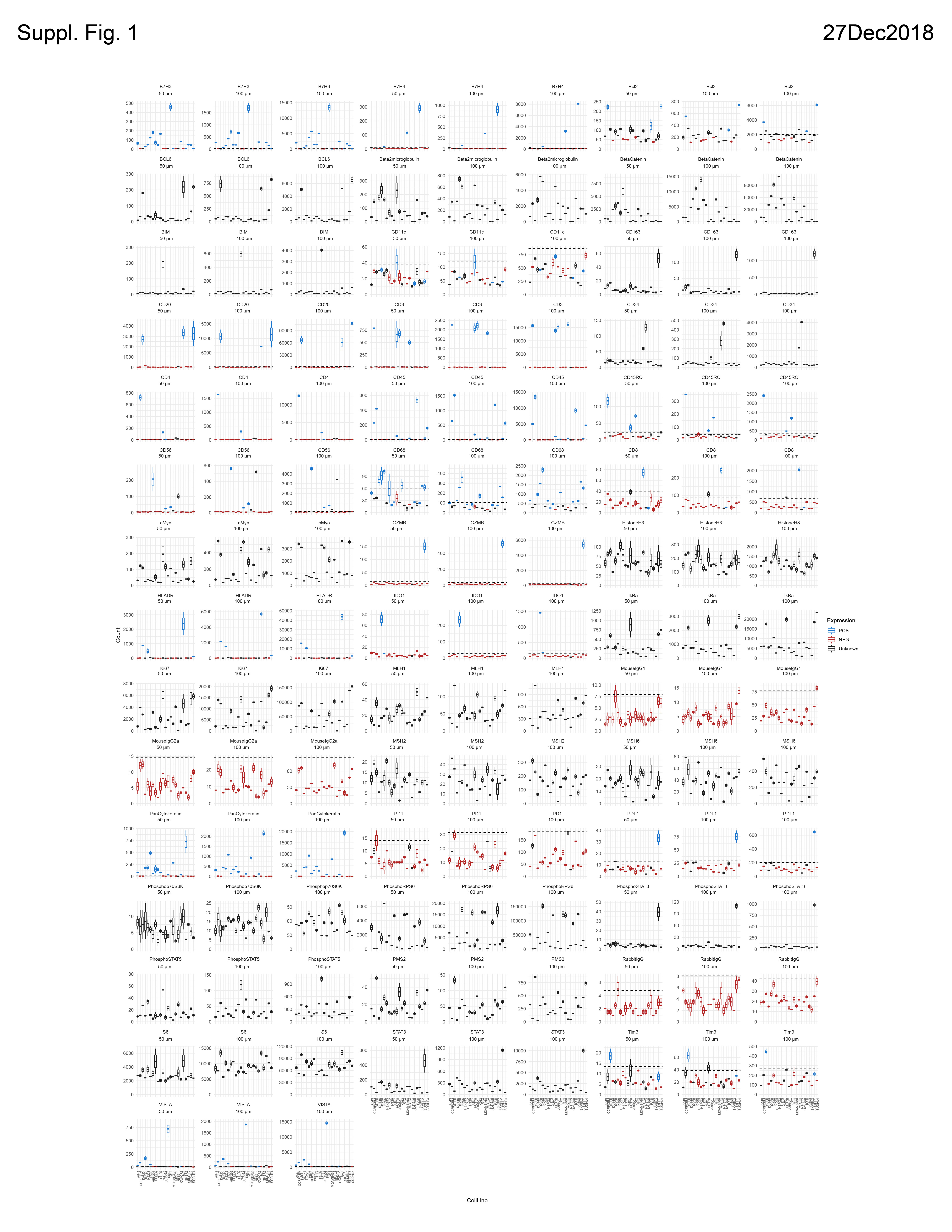

### Supplemental Fig. 2 - hi res

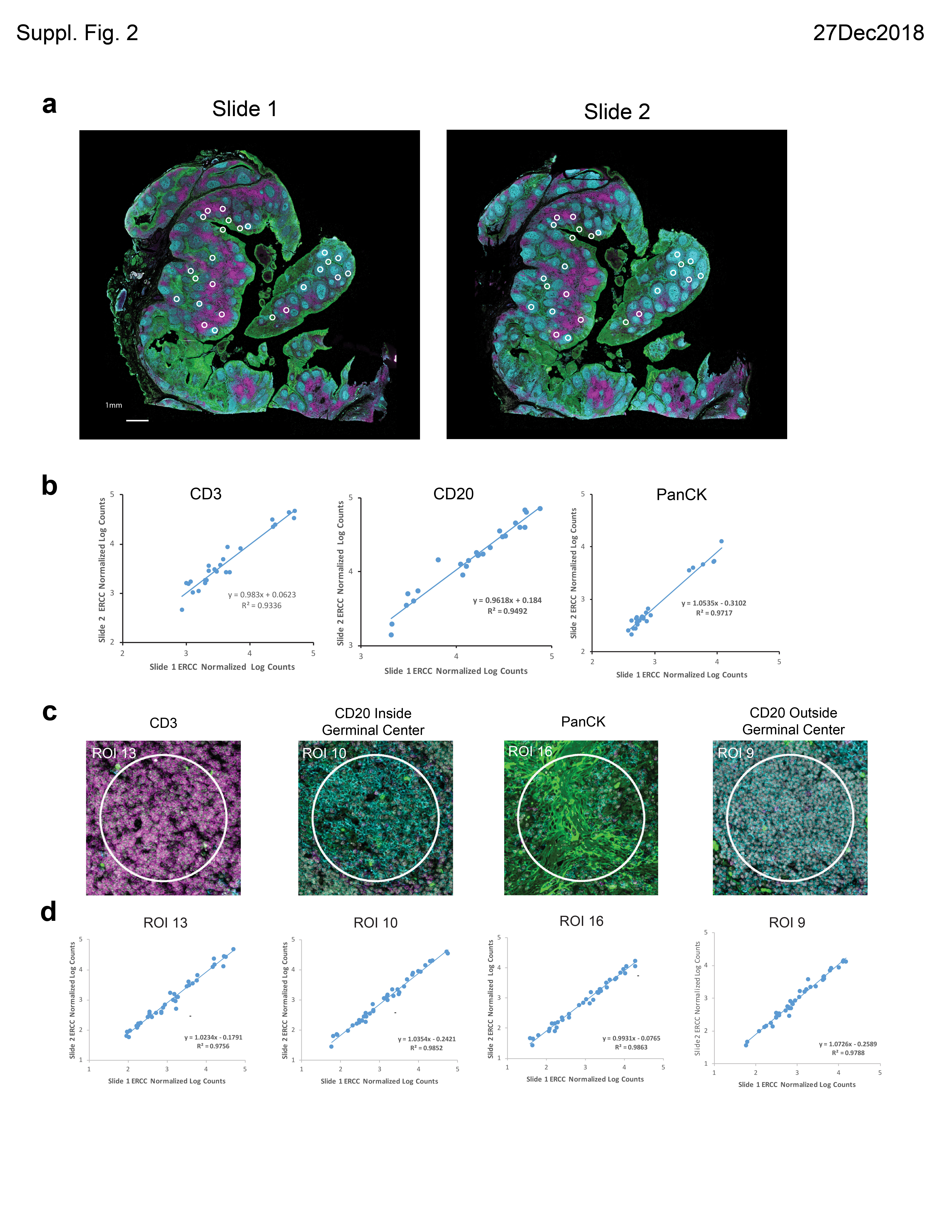

### Supplemental Fig. 3 - hi res

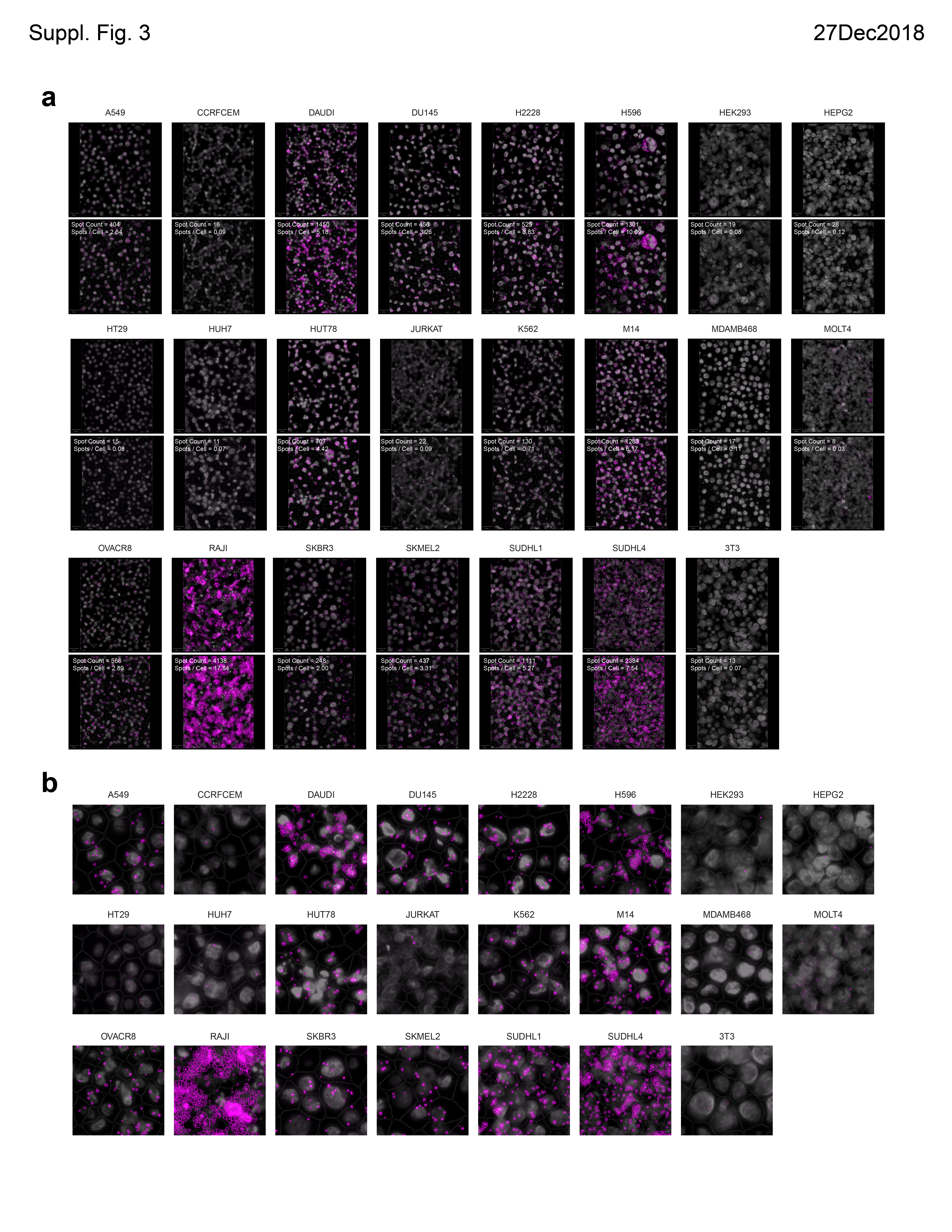

### Supplemental Fig. 4 - hi res

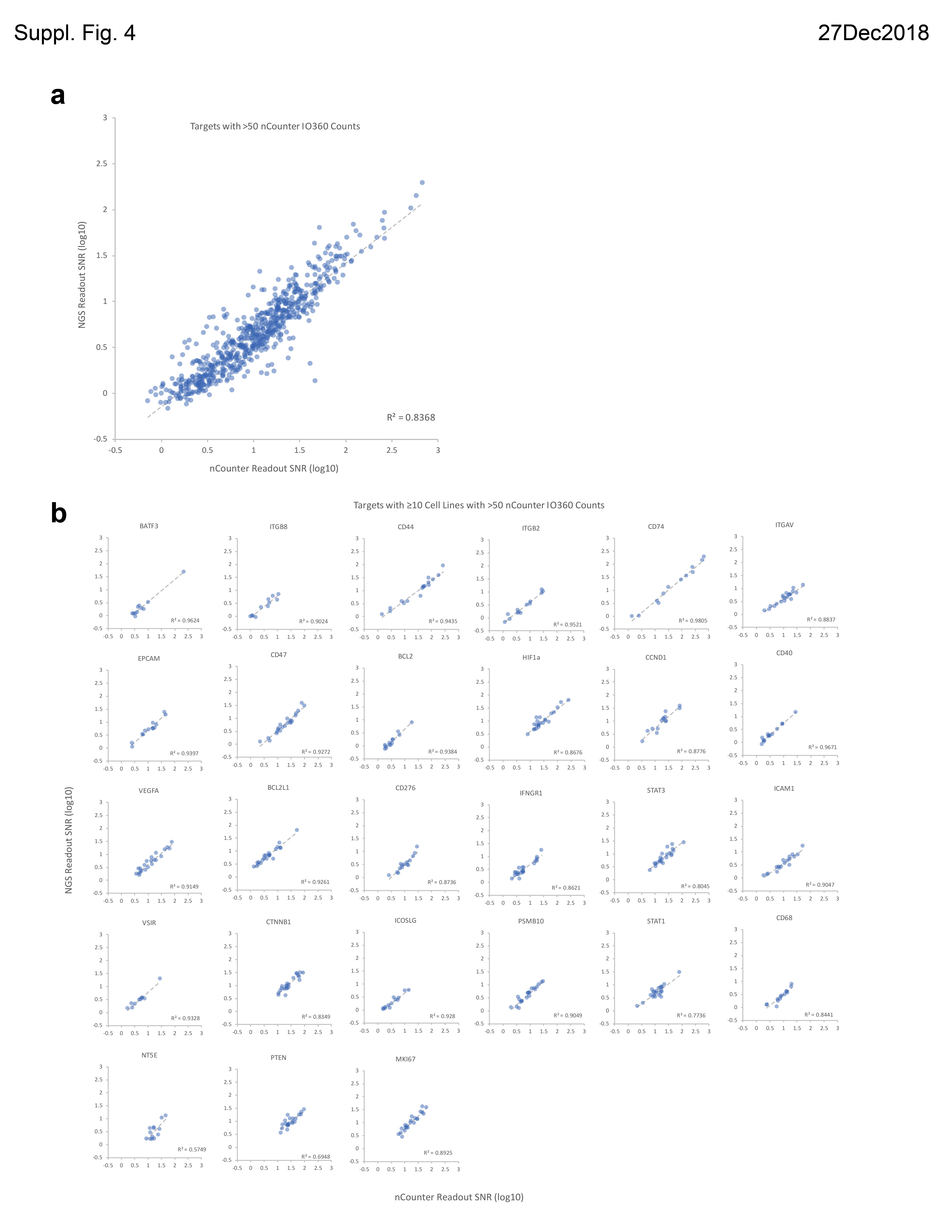

### Supplemental Fig. 5 - hi res

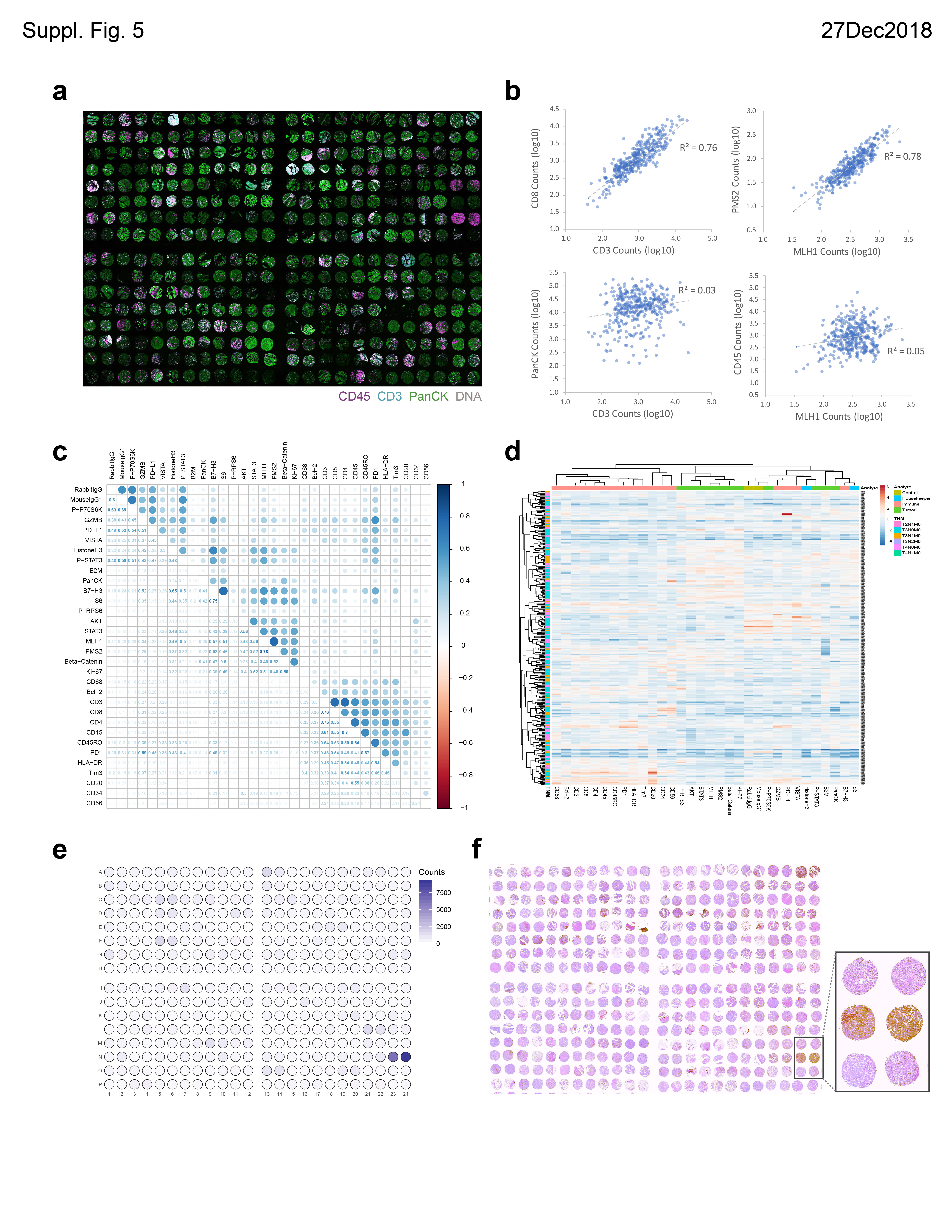

### Supplemental Fig. 6 - hi res

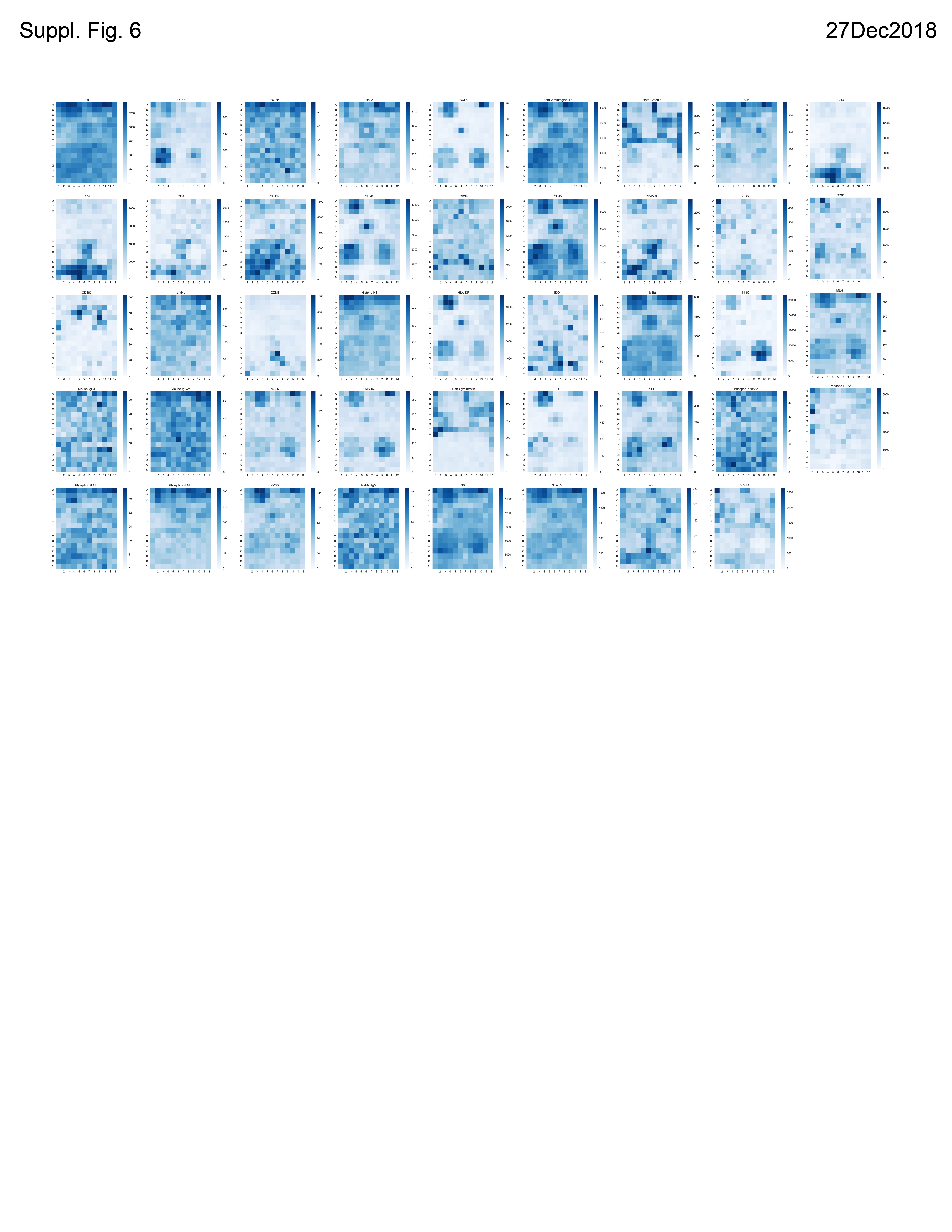
